# Immuno-moodulin is Differentially Expressed in T Cells and Plasma in Obsessive-Compulsive Disorder Patients

**DOI:** 10.1101/2024.12.16.628685

**Authors:** Isobel Alice Blacksell, Matteo Vismara, Christine Maria Lim, Bernardo Dell’Osso, Stefano Pallanti, Eric Hollander, Michele Vendruscolo, Claudio D’Addario, Dianne Cooper, Fulvio D’Acquisto

## Abstract

Immuno-moodulin (Imood), a recently discovered protein expressed in T cells, is associated with anxiety-like behavior in mice. However, its mechanism of action in modulating neuroimmune interactions remains unclear. To investigate this problem, we characterized Imood in human blood and immune cells using neutralizing monoclonal antibodies, revealing its nature as an intrinsically disordered protein (IDP) with unique expression patterns. Our findings indicate that Imood is predominantly expressed intracellularly in peripheral blood mononuclear cells (PBMCs), particularly T lymphocytes, but is absent in polymorphonuclear cells. Upon T-cell activation, Imood exhibits distinct mobilization patterns with increased surface expression. Bioinformatics analysis identified a strong propensity for oligomerization and liquid-liquid phase separation. We also found that T cells from patients with Obsessive-Compulsive-Disorder (OCD) displayed significantly elevated surface Imood expression compared to healthy controls, as well as an altered level of Imood polymerization in the plasma. Taken together, these results elucidate the expression patterns and structural properties of Imood in human immune cells, which open new avenues for OCD diagnostics, and prompt further study for understanding the aetiology of OCD and related disorders.

## Introduction

Obsessive-Compulsive Disorder (OCD) and related conditions are prevalent mental health disorders marked by intrusive thoughts and repetitive behaviors that significantly affect daily functioning (1, 2). Despite their high prevalence (1.9% to 7.8%) (3), the molecular mechanisms underlying these disorders remain unclear (4–6). The complex aetiology of OCD involves interactions between genetic, environmental, and neurobiological factors (5, 7). Although some genetic variations (4) and neurochemical imbalances (8–10) have been identified, a comprehensive understanding of the molecular pathways dysregulated in OCD is lacking. This knowledge gap hinders the development of targeted treatments, as current approaches often rely on broad-spectrum interventions (11–13). Unravelling the molecular intricacies of OCD remains crucial for improving diagnosis, treatment, and prevention strategies.

One intriguing hypothesis concerns the possible role of neuroimmune interactions in promoting OCD onset and progression (14–17). Recent studies have revealed that the nervous system can modulate immune responses through various mechanisms, including the release of neurotransmitters and neuropeptides that interact with immune cells (18, 19). Conversely, the immune system can influence neural function through the production of cytokines and other signaling molecules that affect neuronal activity and behavior (20–22). This two-way relationship has significant implications for the understanding and treatment of a wide range of disorders, including autoimmune diseases, neurological conditions, and mental health disorders. The neuro-immune interaction (23, 24) and continuum (21) hypotheses have opened new avenues for research and potential therapeutic interventions that target the interface between these two systems, potentially leading to more effective treatments for complex disorders that involve both neural and immune components (23, 25–27).

To address these questions, we recently identified a novel modulator of anxiety and repetitive behavior, which we named immuno-moodulin (Imood) (28). First reported as Testis Development-Related Protein (*TDRP*), this gene was initially identified as being predominantly expressed in testis tissue (29, 30). Cloned from a human testis cDNA library, *TDRP* was initially associated with spermatogenesis owing to its high expression in spermatocytes and its developmental regulation during sexual maturation in rats (29). Subsequent studies have revealed a more complex and intriguing biology for this protein.

Contrary to the initial assumptions, *TDRP*-deficient mice showed no significant differences from wild-type littermates in the development of testes, the genitourinary tract, or sperm count (29). Male fertility was not impaired by *TDRP* deficiency alone, suggesting that its function extends beyond reproductive processes. Importantly, the expression of Imood in female subjects (https://www.bgee.org/gene/ENSG00000180190?expression=anat%2Csex) indicated a broader physiological role than that initially hypothesized.

In a previous study (28), we reported *TDRP* as one of the genes differentially expressed in T cells of mice with heightened levels of anxiety-like behavior and increased in peripheral blood mononuclear cells (PBMCs) of OCD patients, providing evidence for the expression of this protein outside the testis. These findings are consistent with a study by the Immunological-Genome-Project-Consortium, which identified 2610019f03Rik/Imood as one of the most differentially expressed genes in T cells compared to B cells, with a fold change value of 84, close to that of the T cell receptor (TCR) chain (31). A further 11 studies (listed in **Table 1**) have identified the gene (*TDRP)* as differentially regulated in various T cell subsets (32–37) as well as during T cell development processes including positive and negative selection (38–41).

**Table 1.** Prior studies identifying differential expression of *TDRP/Imood* (previously known as *C8orf42* or *2610019f03Rik*) in T cell biology. Publications reporting differential expression of *TDRP/Imood* (previously *C8orf42* or *2610019f03Rik*) in T cell populations and during T cell developmental processes, prior to its current nomenclature.

| **Authors** | **Title** | **Journal** | **Reference Number** |
| --- | --- | --- | --- |
| Painter MW *et al.*, | Transcriptomes of the B and T lineages compared by multiplatform microarray profiling | J Immunol | 31 |
| Fu W *et al.*, | A multiply redundant genetic switch 'locks in' the transcriptional signature of regulatory T cells | Nat Immunol | 32 |
| Kaur M *et al.*, | Galectin-3 Regulates gamma-Herpesvirus Specific CD8 T Cell Immunity | iScience | 33 |
| Luckerath K *et al.*, | Immune modulation by Fas ligand reverse signaling: lymphocyte proliferation is attenuated by the intracellular Fas ligand domain | Blood | 34 |
| Joller N *et al.*, | Treg cells expressing the coinhibitory molecule TIGIT selectively inhibit proinflammatory Th1 and Th17 cell responses | Immunity | 35 |
| Lopusna K *et al.*, | Onmtab catalytic activity is critical for its tumor suppressor function in lymphomagenesis and is associated with Met oncogenic signaling | EBioMedicine | 36 |
| Soskic B *et al.*, | Immune disease risk variants regulate gene expression dynamics during CD4(+) T cell activation | Nat Genet | 37 |
| Li P *et al.*, | Reprogramming of T cells to natural killer-like cells upon Bcl1 deletion | Science | 38 |
| Thangavelu G *et al.*, | Programmed death-1 is required for systemic self-tolerance in newly generated T cells during the establishment of immune homeostasis | J Autoimmun | 39 |
| Tsagaratou A *et al.*, | Dissecting the dynamic changes of 5-hydroxymethylcytosine in T-cell development and differentiation | Proc Natl Acad Sci U S A | 40 |
| Ruperez P *et al.*, | Quantitative phosphoproteomic analysis reveals a role for serine and threonine kinases in the cytoskeletal reorganization in early T cell receptor activation in human primary T cells. | Mol Cell Proteomics. | 75 |

In this study, we have extended these analyses by first examining the expression of Imood protein in different types of human immune cells using previously generated monoclonal anti-Imood-neutralizing antibodies (28). Our results revealed that Imood is expressed at high levels in most mononuclear cells, but not in polymorphonuclear cells, and is mobilized from the cytosol to the cell membrane only in activated T cells. We also found that Imood is present in human plasma and exhibits features characteristic of intrinsically disordered proteins (IDPs) (42). T cells from patients with OCD show higher levels of Imood expression and increased polymerization of this protein in the plasma.

Our results suggest that Imood may serve as a molecular link between the immune system and behavioral regulation. These insights lay the groundwork for future investigations into the biological functions of Imood and its potential use in the development of diagnostic and therapeutic tools for OCD and related disorders.

## Materials and Methods

### Reagents

Unless otherwise stated, all chemicals were purchased from Sigma– Aldrich. All antibodies used in the study are reported in **Supplementary Table 1** and in the figure legends where relevant.

### Blood collection

Studies on Imood expression in immune cells of healthy volunteers were performed using freshly isolated blood. Donors provided informed verbal and written consent according to the Declaration of Helsinki for blood collection with prior approval of the procedures by the East London & The City Local Research Ethics Committee (QMERC2019/83). Investigations of Imood expression in patients with OCD and healthy controls were carried out with blood (3.0 ml) collected in Cytomark Transfix vacutainer tubes (43) containing Transfix reagent at a final dilution of 1:10. Eleven OCD outpatients with OCD treated and followed up at the OCD tertiary outpatient Clinic of the “Luigi Sacco” University Hospital in Milan, Italy, were included in the study. Diagnoses were assessed by trained psychiatrists through semi-structured interviews based on DSM-5 criteria (SCID 5 research version, RV). In the cases of psychiatric comorbidity, OCD had to be the primary disorder, causing the most significant distress and dysfunction and providing the primary motivation to seek treatment. Patients were excluded from the study if they had recent or current alcohol or substance abuse (last 3 months), as well as other medical conditions, including autoimmune diseases, owing to their potential influence on gene expression. For the same reason, the lifetime history of trauma (according to DSM-5), as well as the current presence of relevant psychological stress, were considered exclusion criteria. Clinical assessment included the collection of the following demographic and clinical variables: sex, age, age at onset, and current pharmacological treatment. In addition, illness severity was measured using the Yale-Brown Obsessive-Compulsive Scale (44). Patients who had maintained their pharmacological treatment for at least one month were enrolled in the study. A convenience sample of healthy controls (recruited from study investigators, their relatives, and acquaintances) was included as the control condition. Control subjects (n=14) were volunteers matched for sex, age, and ethnicity, with no psychiatric diagnosis as determined by the SCID-5, and no family history of major psychiatric disorders in first-degree relatives (as assessed by the Family Interview for Genetic Studies). Blood was collected from fasting donors between 2 pm and 4 pm. All participants provided written informed consent to participate in the study, which included the use of personal and clinical data. This study was approved by the Ethics Committee of Milan Area 1 (No. 2021/ST/165).

### Peripheral blood mononuclear cell (PBMCs) and polymorphonuclear (PMN) leukocyte isolation and activation

Fresh venous blood from healthy volunteers was drawn into 3.2% sodium citrate and diluted 1:1 in RPMI 1640 (Thermo Fisher; 11835030) before separation through a double density gradient as previously described. Briefly, the cells were resuspended in incomplete RPMI 1640 medium, counted prior to resuspension in complete RPMI-1640, plated, and used for in vitro experiments. Pan-PBMC stimulation was achieved by incubating cells (3x10^5^ cells) with a cell stimulation cocktail (00-4970-03; eBioscience) containing phorbol-12-myristate-13-acetate (PMA) (80 nM) and ionomycin (1.34 μM) for 4 h at 37 °C. To selectively activate T cells, PBMCs were seeded in 48-well plates coated with plate-bound anti-CD3 (5ug/mL) (clone OKT3; BioLegend). Anti-CD28 (clone 28.2; BioLegend) was added to the culture at a concentration of (5 mg/mL) and the cells were incubated overnight at 37 °C. Both PMA- and anti-CD3/CD28 stimulated cells were stained for surface and intracellular Imood expression, as described below.

### Flow cytometric analysis of Imood expression

For Imood surface staining, isolated PBMCs were resuspended in 200 µL of FACS buffer (DPBS^-/-^) containing 0.02% bovine serum albumin (BSA; Sigma), mixed 1:1 with a solution containing blocking human immunoglobulins (hIgGs) (Sigma, G4386) at 160 µg/mL in DPBS^-/-^ and incubated on ice for 10 min. For whole blood staining, 50 µL of blocking hIgG was added to 50 µL of whole blood for 10 min at RT. All isolated cell staining was completed on ice, and whole-blood staining was completed at RT. Thereafter, the cells were washed and resuspended in FACS buffer containing antibodies for phenotypic surface markers as follows: for PBMCs, CD45, CD3, CD4, CD8, CD14, CD16, CD19, and CD56 were used; for PMN, CD66b, CD45, CD11b, and for whole blood CD45, CD4, and CD8 were used. Master-mixes additionally contained commercially available polyclonal rabbit anti-Imood antibody (Novus Biologicals; NBP1-93675), 1B10 or 1C4 antibodies (both at 40 µg/mL) or respective isotype controls. Samples were washed and incubated with AF488-conjugated secondary antibody (10 µg/mL). After labelling, isolated cells were fixed with 1% PFA and stored prior to analysis using a BD LSR Fortessa flow cytometer. Whole blood samples were fixed with 1x fixative of the FixPerm eBioscience kit and stored prior to analysis on an Attune NxT flow cytometer (Thermo Fisher). For intracellular staining, cells were processed using a FixPerm kit (eBioscience; 00-5523-00) according to manufacturer’s instructions, followed by analysis on an Attune NxT flow cytometer. Flow cytometry data were analyzed using FlowJo 10.7.1 (FlowJo LLC).

### Immunocytochemistry

Twelve millimetre-round coverslips, #1 thickness (Fisher Scientific), were coated with 1% Alcian Blue (Sigma; 33864-99-2) solution. Isolated PBMCs were diluted to a suspension of approximately 1x10^6^ in serum-free media and 100 μL was added per coverslip. Cells were allowed to adhere at 37 °C for 30 min and then fixed in 4% PFA. Coverslips were blocked for 1 h at RT with 3% goat serum (Sigma; G9023) in DPBS^-/-^ alone for membrane staining and with 0.1% Triton-X (Sigma; X100) to permeabilize the cells for intracellular staining. Cells were incubated with 100 μL of primary antibody diluted in 3% goat serum in PBS -/-with or without 0.1% Triton-X for 1 h at RT. After the appropriate secondary antibody was added for 1 h at RT. Coverslips were mounted onto superfrost slides (Fisher Scientific) using a drop of Fluoroshield™ with DAPI (Sigma; F6057). Slides were stored at 4°C in the dark until visualization. Cells were visualised at 63x magnification on a Zeiss LSM-800 confocal microscope, and images were captured using Zeiss Zen-Blue software. Images were processed using Fiji (ImageJ)(45) .

### Western blotting analysis

Plasma (5 µL) was diluted in 95 µL of µL 1x lithium dodecyl sulfate (LDS) Laemmli buffer (Invitrogen: NP007), diluted 4x with dH_2_O, containing 1 mM DTT (ThermoFisher; 707265), and boiled at 95 °C for 5 min. Samples (supernatants/serum/lysates) were subjected to electrophoresis on an SDS-10% polyacrylamide gel (BioRad; 4561033) followed by transfer onto polyvinylidene difluoride (PVDF) Immobilon-P Transfer membranes (Millipore; IPVH00010). Non-specific binding was blocked for 1 h at RT with 5% non-fat dry milk (Marvel). Primary antibodies diluted in 5% milk in DPBS-T were incubated overnight at 4 °C. Following washing secondary antibodies conjugated to horseradish peroxidase were applied for 1 h at RT. Immunoblotting and visualization of proteins was performed by enhanced chemiluminescence (ECL; Millipore; WBLUF0500) according to the manufacturer’s instructions. Membranes were visualized using an Azure 400 Biosystem.

### Statistics

All data were subjected to normality testing using the Shapiro-Wilk test. Data are displayed as mean ± SEM. According to the nature of the data obtained and the normality test results, parametric data were analyzed using a t-test (2-tailed) or paired t-test (2-tailed). For non-parametric data, either the Mann-Whitney test or Wilcoxon matched-pairs signed rank test was performed. All statistical analyses were performed using GraphPad PRISM software v8.0-9.1.2.

## Results

### Imood expression in human blood

We began our investigation of Imood expression in human immune cells by comparing its surface and intracellular expression in two main classes of blood leukocytes: PMN and PBMC (**Figure 1A**). We employed a commercially available polyclonal rabbit anti-Imood antibody and two monoclonal neutralizing anti-Imood antibodies (1B10 and 1C4; (28)) together with the pan-leukocyte marker CD45.

**Figure 1:**
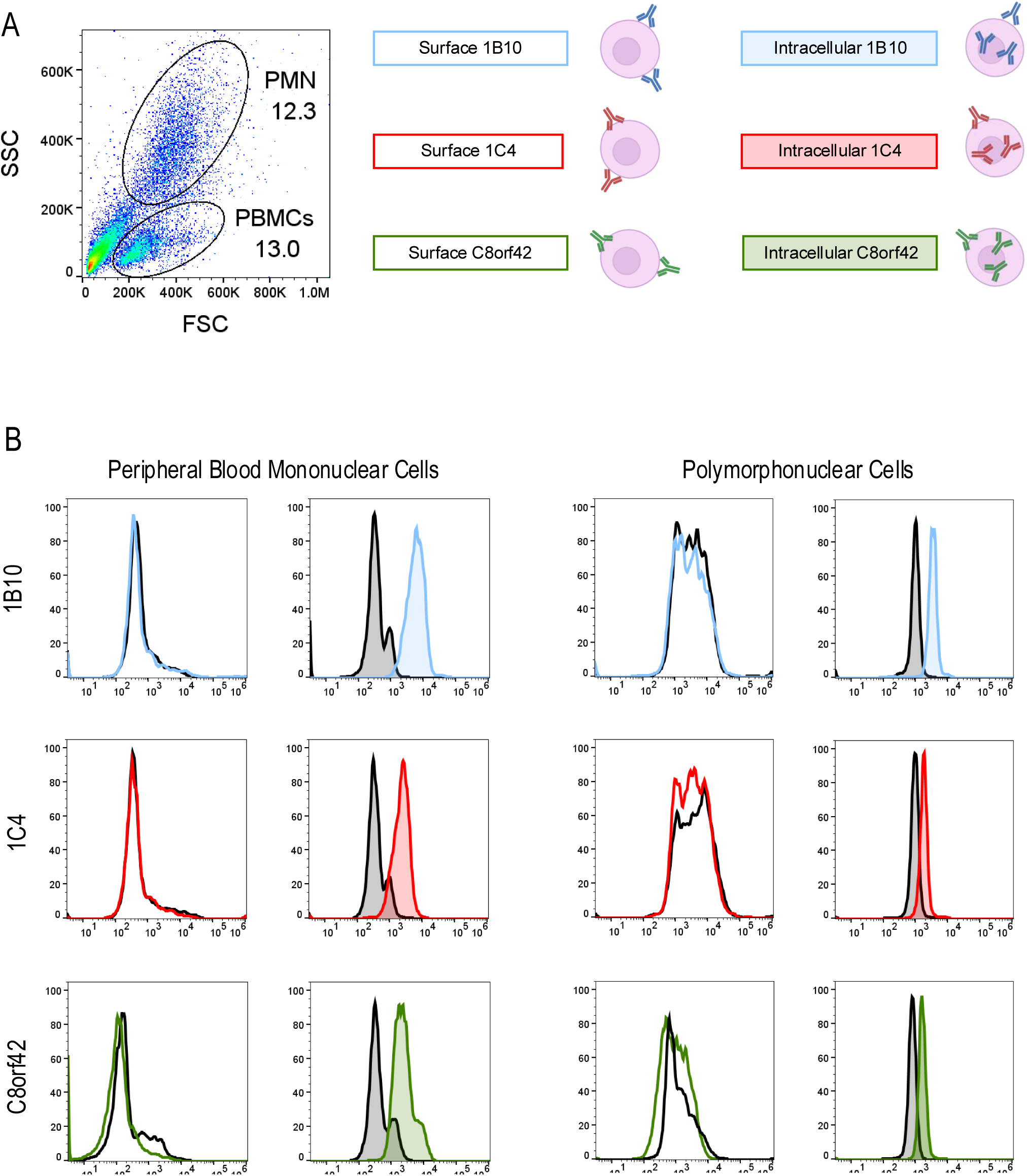
Expression of Imood in human immune cells under basal conditions. **(A)** Flow cytometry gating strategy for identifying polymorphonuclear neutrophils (PMN) and peripheral blood mononuclear cells (PBMC) in whole blood based on forward scatter (FSC) and side scatter (SSC) properties. The percentages of PMN (12.3%) and PBMC (13.0%) are indicated. (**B**) Comparative analysis of surface and intracellular Imood expression in PBMC (left panel) and PMN (right panel) using three different anti-Imood antibodies: in-house mAb 1B10 (blue), 1C4 (red) and commercial C8orf42 (green). Surface expression is represented by empty histograms, while intracellular staining is shown with filled histograms. Gray histograms indicate isotype controls. The histograms are from a single subject and representative of n=3-5 healthy volunteers with similar results.

Imood was not detectable on the surface of either PMN or PBMC with any of the three anti-Imood antibodies tested (**Figure 1B**). However, the analysis of intracellular Imood levels yielded different results. PMNs showed very low expression of Imood intracellularly with all three antibodies, whereas PBMC demonstrated a significant increase in median fluorescence intensity (MFI; data not shown). The 1B10 antibody provided the highest increase (approximately 17-fold) over the isotype control, compared to 1C4 and polyclonal anti-Imood (both increased approximately 6-fold) (**Figure 1B**). These results suggest that under healthy physiological conditions, Imood is not expressed on the cell surface, but is primarily stored in the intracellular compartments of mononuclear cells and is expressed at very low or undetectable levels in innate immune cells, such as granulocytes.

As several repository platforms for the collection of functional information on proteins report Imood to be a secreted protein present in the blood (https://db.systemsbiology.net/sbeams/cgi/PeptideAtlas/GetProtein) (46, 47), we tested Imood levels in plasma using the same antibodies used for flow cytometry. Immunoblot of as little as 0.5-1.0 μL of plasma provided a positive signal with all three antibodies (**Figure 2A**). Both 1B10 and 1C4 immunoblots showed the expected Imood band at approximately 21 kDa, together with other higher molecular weight bands corresponding to dimers (42 kDa), trimers (63 kDa), and polymers (>100 kDa). Conversely, the polyclonal anti-Imood antibody revealed two bands: one at approximately 15 kDa and the second at approximately 130 kDa, but no or faint monomers, dimers, or trimers (**Figure 2A**). We explained this difference as the antibody was generated against a deletion mutant of Imood (amino acids 35-185) that lacked the disordered N-terminus (**Figure 2D**), which likely plays a role in Imood oligomerization. In addition, recombinant Imood used as a positive control in all blots showed no or a faint signal with 1B10, as previously reported, and a strong detectable band at approximately 27 kDa (21 kDa of full-length Imood plus 3 kDa of c-Myc tag on the C-terminus) with both 1C4 and polyclonal anti-Imood (**Figure 2A**).

**Figure 2:**
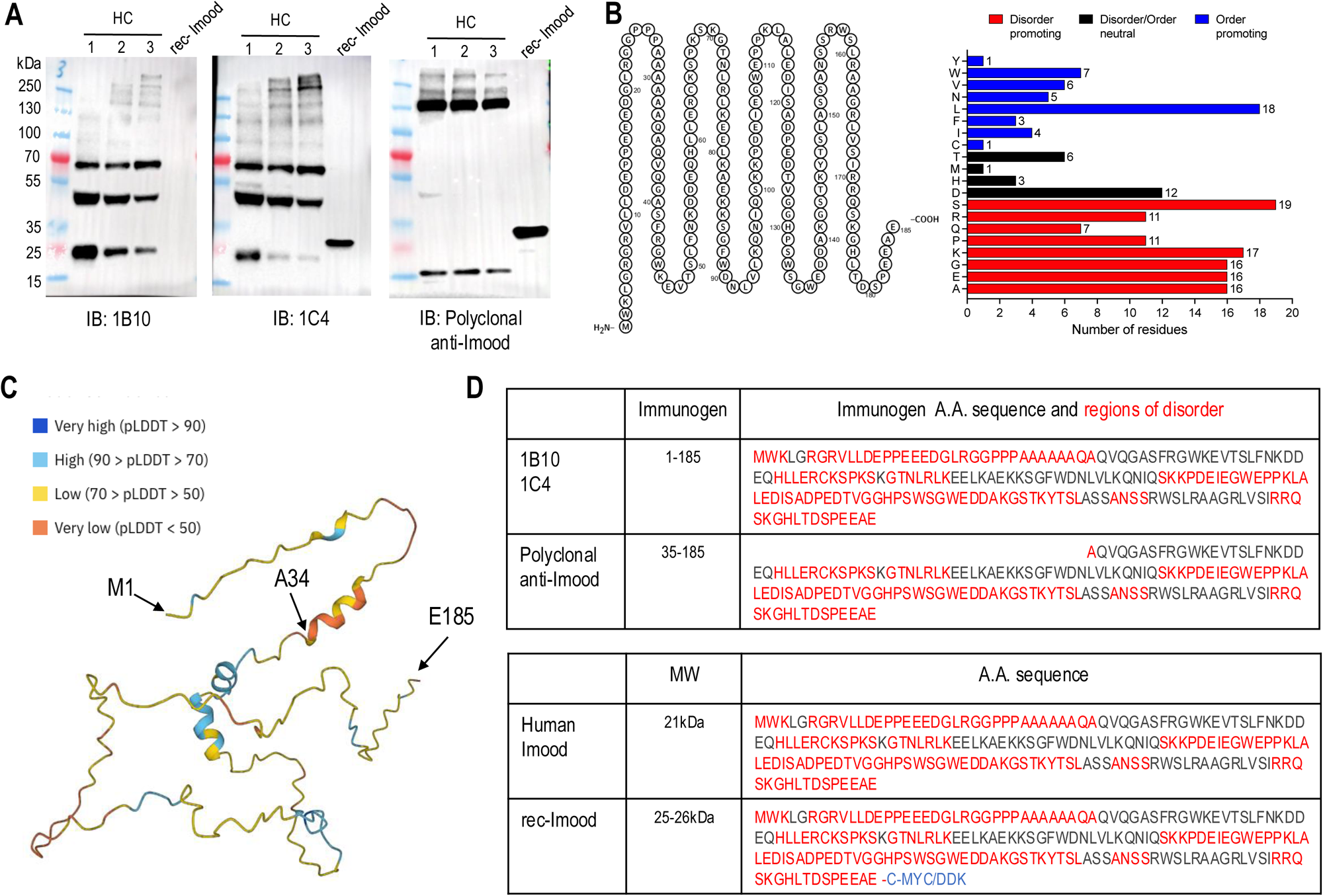
Characterization of Imood expression in plasma and analysis of its structural properties. (**A**) Western blot analysis of human plasma (0.5-1.0 μL) using three different anti-Imood antibodies: 1B10, 1C4, and polyclonal anti-Imood. HC lanes (1–3) represent human plasma samples from different individuals. Recombinant Imood (rec-Imood) is used as a positive control. 1B10 and 1C4 antibodies detect multiple bands corresponding to monomeric (21 kDa), dimeric (42 kDa), trimeric (63 kDa), and polymeric (>100 kDa) forms of Imood. The polyclonal antibody primarily detects two bands at ∼15 kDa and ∼130 kDa. Rec-Imood shows a strong band at ∼27 kDa with 1C4 and polyclonal antibodies, but weak or no signal with 1B10. (**B**) Amino acid composition analysis of Imood. The bar graph shows the distribution of disorder-promoting (red), disorder-neutral (black), and order-promoting (blue) amino acids in the Imood sequence. Numbers indicate the count of each amino acid type. (**C**) 3D structure prediction of Imood using AlphaFold. The colour scheme represents the predicted local distance difference test (pLDDT) scores: very high (dark blue, >90), high (light blue, >70), low (yellow, >50), and very low (orange, <50). The model shows predominantly disordered regions with a few short, ordered segments. (**D**) Table summarizing the epitope regions recognized by different anti-Imood antibodies (1B10, 1C4, and polyclonal) and the amino acid sequences of human Imood and recombinant Imood. Regions of disorder are highlighted in red.

To better understand the nature of the different Imood bands detected, we searched public protein databases (48, 49) for known larger precursors or polymerization domains within the Imood amino acid (AA) sequence (**Figure 2B**). This showed the presence of several phosphorylation/mono-methylation sites (K3, S41, S66, K145, Y146, S173) as well as a high abundance (113 out of 185 in total) of disorder-promoting AA (50), such as A (n=16), E (n=19), G (n=16), K (n=17), P (n=11), Q (n=7), R (n=11), and S (n=19), compared to the number of order/disorder-neutral (D, H, M, and T) or order-promoting (C, I, F, L, N, V, W, Y) residues, accounting for approximately 60% of the protein sequence. Consistent with these findings, the prediction of Imood structure by AlphaFold2 (51) showed that apart from a few short-ordered regions with a high predicted local distance difference test (pLDDT; between 70 and 90) (52), most of the proteins showed a low (between 70 and 50) or very low (below 50) pLDDT, indicative of the presence of intrinsically disordered regions (IDRs) (53) (**Figure 2C**).

To further substantiate the presence of IDRs in Imood, we sought to quantify the intrinsic disorder on a per-residue basis using the RIDAO suite (https://ridao.app/), which encompasses the analysis of the most known software predictors of disorders (PONDR® VLXT (54), PONDR® VL3 (55), PONDR® VSL2 (56), IUPred-Long, and IUPred-Short (57)). These platforms classify proteins based on their percentage of predicted disordered residues (PPDR scores), where proteins are considered highly ordered (PPDRD<D10%), moderately disordered (10%D≤DPPDRD<D30%), and highly disordered (PPDRD≥D30%). The results for Imood are shown in **Supplementary Figure 1A** and the corresponding data are 59% for PONDR® VLXT (i.e., highly disordered), 100% for PONDR® VL3 (i.e., highly disordered), 100% for PONDR® VSL2 (i.e., highly disordered), 21% for PONDR-FIT (i.e., moderately disordered), 100% for IUPred-Long, and IUPred-Short (i.e., highly disordered).

To assess the functionality of intrinsic disorder in Imood, we used the D^2^P^2^ platform (58) (https://d2p2.pro/), which provides a correlation between post-translational modifications (PTMs) and disorder-based binding potential. The Imood D^2^P^2^ output demonstrated relative agreement between the nine disorder-based predictors, where scattered regions of disorder predicted throughout the protein were present (**Supplementary Figure 1B,C**). The regions of Imood that demonstrated the highest agreement among the nine per-residue disorder predictors were around the N-terminus (amino acids 1-40) and C-terminus (amino acids 170-185). Analysis of molecular recognition feature (MoRF) regions – domains that interact with structured partner proteins also revealed five distinct regions (1-10; 28-53; 86-96; 147-153; 156-170). Three putative phosphorylation sites (Ser 41, Ser 66, and Ser 173) were also identified.

Many IDPs condense into liquid-like droplets, that is, a biomolecule-rich phase coexisting with a more dilute solution (59, 60). This process is known as liquid–liquid phase separation (LLPS) and is one of the ways in which cells compartmentalize proteins (61–63). LLPS plays a crucial role in many diseases (64–67), as its dysregulation leads to the maturation of biomolecular condensates into hydrogel-like assemblies, promoting the formation of toxic oligomers and fibrils. The propensity of a given protein to undergo LLPS can be evaluated computationally using FuzDrop (68). One advantage of this tool is its ability to group proteins based on their LLPS propensity (p_LLPS_). Proteins capable of spontaneous LLPS are classified as droplet-driving proteins, whereas those that require additional interactions to form droplets are classified as clients. As per FuzDrop, LLPS drivers are proteins with an overall p_LLPS_ D≥D0.60, whereas proteins with lower overall pLLPS but containing droplet-promoting regions (DPRs) (69), defined as consecutive residues with pLLPS ≥D0.60 will likely serve as droplet clients. **Supplementary Figure 2A** shows that Imood features a high p_LLPS_ score (0.85) and contains multiple droplet-promoting regions (1-26; 117-144; 172-185) as well as aggregation spots (9-19; 101-110; 124-130; 136-144; 162-167) i.e. residues and regions that may promote the conversion of the liquid-like condensed state into a solid-like amyloid state. These predictions suggest that Imood forms droplets and amyloid-like fibers.

The predictions provided by FuzDrop include the probability of a protein having residues and regions with cellular context-dependence (i.e., regions containing residues characterized by the ability to switch between disordered and order-based binding). **Supplementary Figure 3A** shows eight regions with context-dependent interactions (70) (residues 9–19, 23–29, 69–75, 85–90, 96–118; 124-130; 136-145; and 157-176). These regions overlap with, are included in, or are in close proximity to the eight IDPRs with 75% agreement within different predictors found in Imood (e.g., residues 1-3; 6-35; 59-69; 71-77; 100-141;144-149; 153-156; 170-185). Furthermore, only one of the eight IDPRs with context-dependent interactions also includes a phosphorylation site (Ser173), suggesting that the functionality of this region can be regulated by PTMs, whereas the other two putative PTMs are either outside (Ser41) or within (Ser66) of these regions.

Collectively, these bioinformatic analyses suggest that Imood is an IDP with a high propensity for liquid-liquid phase separation, potentially forming functional condensates or aggregates in cellular contexts. To further explore this possibility, we investigated the expression patterns of Imood across the different subpopulations of PBMCs that were found to express the highest levels of this protein.

### Assessment of Imood Expression in Peripheral Blood Mononuclear Cells

We measured Imood expression in the main subclasses of PBMCs, T and B cells, monocytes, and natural killer (NK) cells using classical phenotypic markers (CD4 and CD8 for T cells, CD16 for monocytes, CD19 for B cells, and CD56 for NK cells). The gating strategy employed for these populations is shown in **Figure 3A**. **Figure 3B** presents the corresponding histograms for surface and intracellular Imood staining.

**Figure 3:**
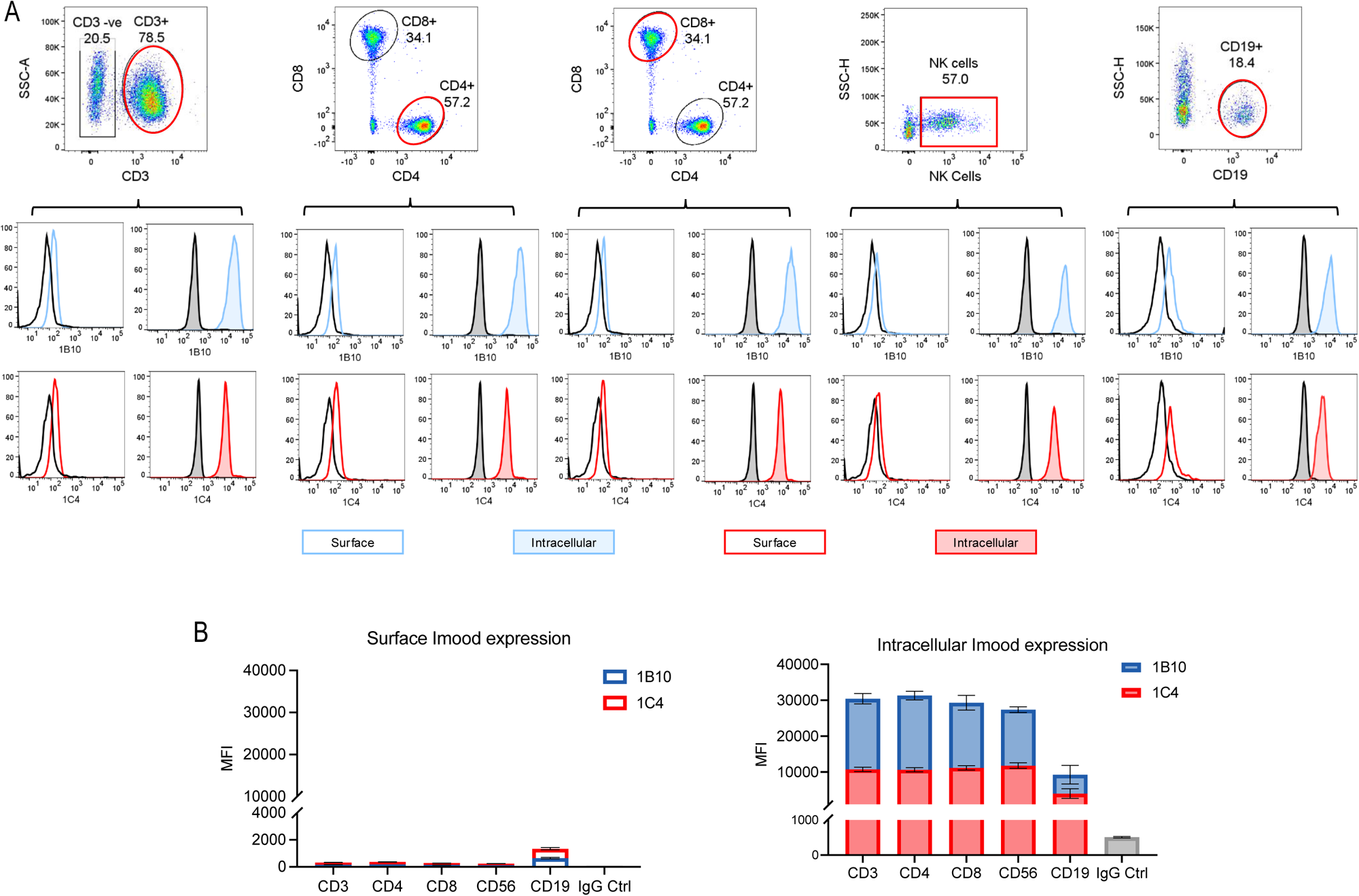
Analysis of Imood expression in mononuclear cell subpopulations. (A) The top row shows the gating strategy for identifying different mononuclear cell subpopulations: CD3+ T cells (further divided into CD4+ and CD8+ subsets), CD16+ monocytes, CD19+ B cells, and CD56+ NK cells. Each subpopulation is highlighted with a coloured gate in the respective flow cytometry plot. The bottom two rows display histograms for Imood staining in each cell subpopulation. Histograms show Imood surface staining (unfilled histograms) and intracellular staining profiles (tinted histograms) using mAbs 1B10 (blue) and 1C4 (red). IgG control is shown in black. Cumulative stacked bar graphs show MFI values for surface (left) and intracellular (right) Imood expression in T cells (CD3+, CD4+, CD8+), CD56+ and CD19+ positive cells. Blue bars represent staining with the 1B10 antibody, and red bars represent staining with the 1C4 antibody. Results are expressed as mean ± SEM (n=3 biological replicates).

This analysis revealed low surface staining for both 1B10 and 1C4 antibodies in all cell subtypes, as revealed by an almost overlap with the staining obtained with the isotype control (**Figure 3A**, empty histograms and **3B**, left panel). Conversely, intracellular staining showed a marked difference in MFI compared to the isotype control and similar levels between T cells and NK cells, but not B cells that expressed approximately a third less. In addition, staining with the 1B10 antibody was uniformly higher than that with 1C4 across the evaluated cell types (**Figure 3A**, filled histograms and **3B**, right panel).

A similar analysis of primary classes of circulating innate blood cells, including CD14+ monocytes, CD16+ monocytes, and CD66B+ neutrophils, corroborated our previous findings (**Figure 1**), showing that neutrophils lack detectable Imood expression at both the surface and intracellular levels (**Supplementary Figure 4A**, **B**). In contrast, classical CD14+ monocytes demonstrated appreciable Imood membrane expression, whereas non-classical CD16+ monocytes did not exhibit a discernible difference in Imood staining compared with the IgG control (**Supplementary** Figure 4A,B, left panel). Intriguingly, intracellular staining comparison between these monocyte subsets revealed an inverse relationship, with CD16+ monocytes expressing Imood at levels 2-fold to 3-fold higher than their CD14+ counterparts (**Supplementary** Figure 4A,B, right panel).

### Differential Mobilization of Imood in Activated T Lymphocytes

After analyzing the patterns of Imood expression in all immune cells at basal levels, subsequent experiments were performed to determine whether Imood expression and localization were altered by cellular activation. To this end, PBMCs were first stimulated with the pan-leukocyte activator PMA (71) so that all cells could be stimulated simultaneously. No significant modifications in Imood expression were observed on the surface or intracellularly in activated CD14+ monocytes or CD19+ B cells compared with their resting states (**Supplementary Figure 5**). We were unable to stain NK cells in the same samples because of their reduced viability (data not shown).

Conversely, activation elicited a distinct expression pattern in T lymphocytes, with notable differences between the 1B10 and 1C4 staining. Following 1B10 staining, both CD4+ and CD8+ T cells exhibited minimal changes in the percentage of Imood+ cells post-activation (**Figure 4A**).

**Figure 4.**
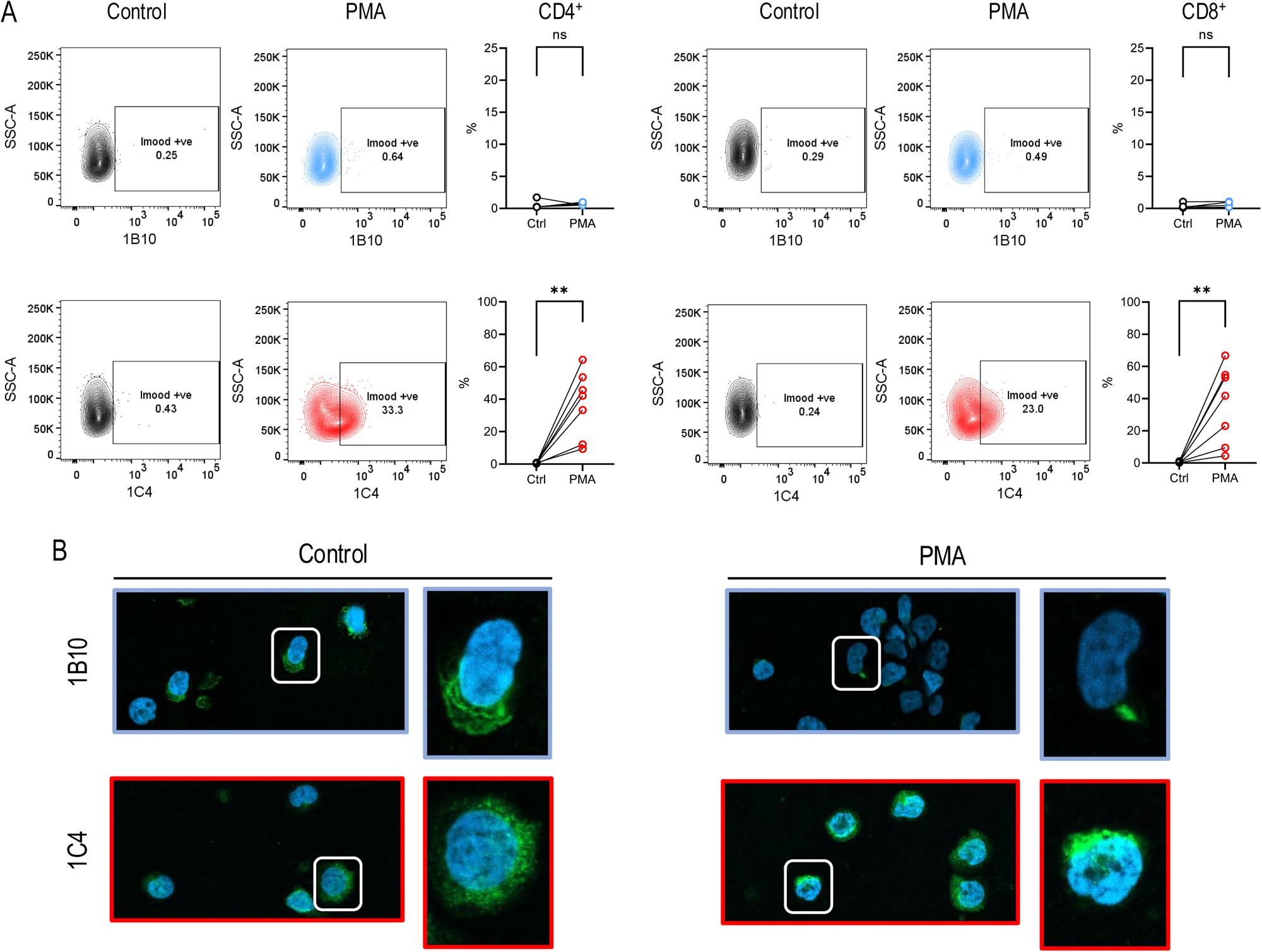
1C4-detected Imood is significantly upregulated to the surface of CD4+ and CD8+ T cells upon stimulation of PBMCs. PBMCs were stimulated with PMA (80 nM) and ionomycin (1.3 μM) for 4 h prior to staining for flow cytometry or immunofluorescence imaging using confocal microscopy. (**A**) Imood-positive gates were drawn on the right-hand side of the CD4 or CD8 control populations to observe the % right-hand shift into the Imood positive gate. Gates were set for each individual control and matched to their comparative intra-donor-stimulated counterparts to observe increased protein expression. Graphs display individual donor-dependent percentages of entire gated CD4+ and CD8+ populations matched for intra-donor changes between the control and stimulus groups. Normal data distribution was assessed using the Shapiro-Wilk test. Results are expressed as mean ± SEM (n=7 biological replicates) using a paired t-test or Wilcoxon matched-pairs signed-rank test: *p< 0.05; **p< 0.01. (**B**) Illustrative confocal microscopy images visually confirming the alterations in protein localization observed by flow cytometry. Imood expression was detected using anti-rat AF488 (green) following incubation with 1B10 and 1C4 rat monoclonal primary antibodies, and cell nuclei using DAPI (blue).

In contrast, 1C4 staining showed a dramatically different trend, with a substantial increase in the percentage of Imood+ cells. For CD4+ T cells, the percentage of 1C4+ cells increased from approximately 1% in the control to approximately 40% in PMA-activated cells. Similarly, for CD8+ T cells, the percentage of 1C4+ cells increased from approximately 1% in the control to approximately 30% in PMA-activated cells (**Figure 4A**).

The intracellular Imood levels mirrored the changes observed at the cell surface with either antibody (data not shown). These results suggest that while the overall surface expression of Imood (as detected by 1B10) may decrease slightly upon activation, there is a significant increase in a specific epitope or conformation of Imood recognized by 1C4, potentially indicating a functional change in the protein upon T-cell activation.

Fluorescence microscopy confirmed the flow cytometry findings for surface staining, with control T cells displaying a faint and polarized Imood staining pattern with 1B10, which was markedly diminished or absent upon PMA stimulation. Meanwhile, 1C4 staining, which was more diffuse and punctate in resting cells, was more pronounced in activated T-cells (**Figure 4B**). Collectively, these observations suggest the presence of discrete Imood reservoirs within T cells that are differentially mobilized upon activation.

To dissect the dynamics of Imood expression upon T-cell activation, we stimulated the cells with anti-CD3 and anti-CD28 antibodies (72–74), which closely mimic physiological T-cell receptor (TCR) engagement. This approach yielded outcomes with distinct characteristics compared with PMA stimulation. Upon activation, the percentage of Imood+ cells, detected by 1B10 staining, decreased in both CD4+ and CD8+ T cells by approximately 8% and 16%, respectively (**Figure 5**). This decrease was statistically significant for CD8+ T cells (P ≤ 0.05). Conversely, the percentage of Imood+ cells detected by 1C4 staining increased markedly, with CD4+ T cells increasing by 45% and CD8+ T cells by 47%, both reaching statistical significance (P ≤ 0.01 for CD4+ and P ≤ 0.001 for CD8+ T cells).

**Figure 5:**
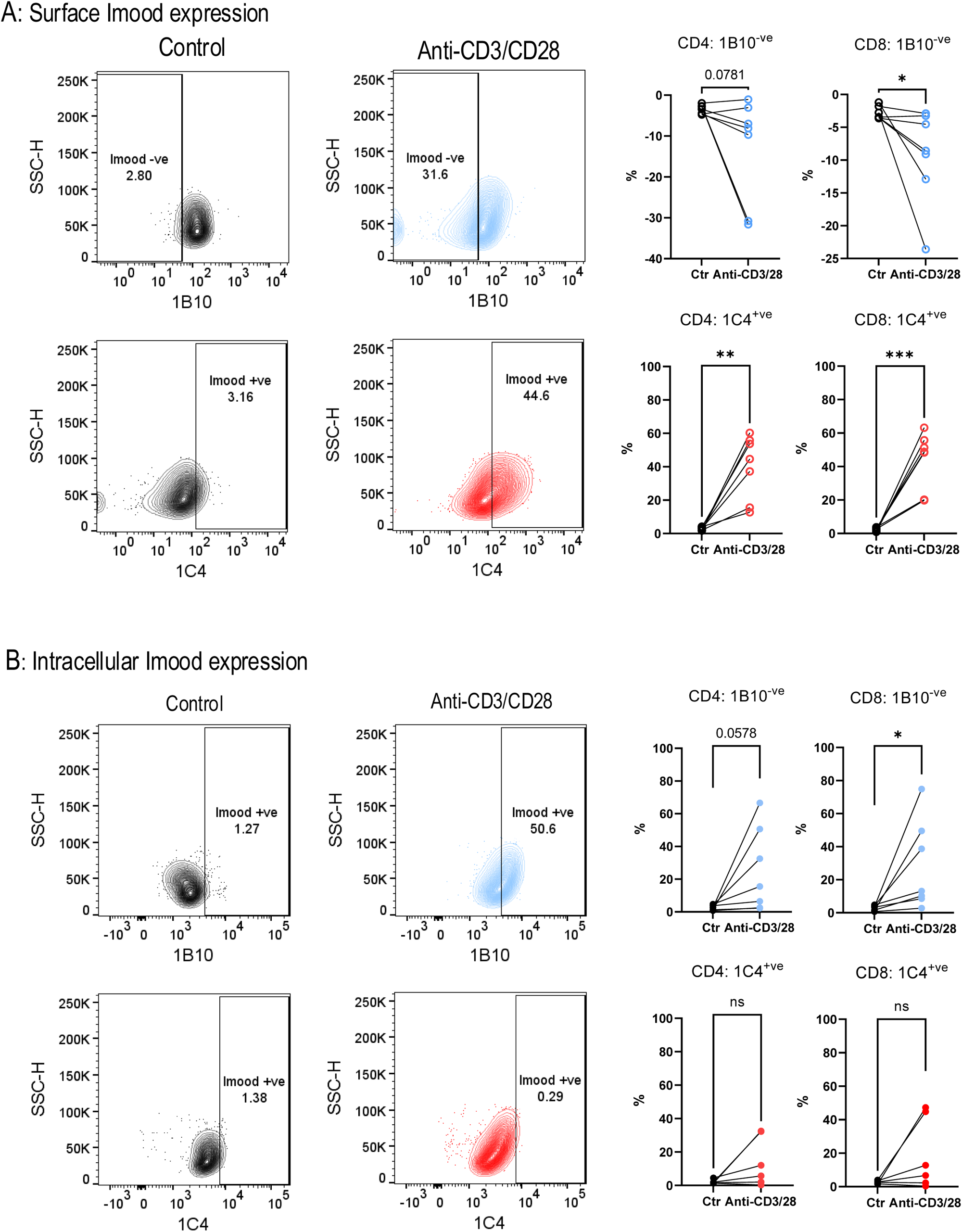
Analysis of Surface and Intracellular Imood Expression in CD4+ and CD8+ T Cells. (**A**) Surface Imood Expression: Flow cytometry analysis was performed to evaluate surface Imood expression on CD4+ and CD8+ T cells, comparing unstimulated controls with cells activated via anti-CD3/CD28. Imood expression was detected using two antibodies, 1B10 and 1C4. Representative contour plots show the percentage of Imood-positive cells, with 1B10 (blue) and 1C4 (red) staining plotted on the X-axis against side scatter height (SSC-H). Paired before-after graphs (right) depict the percentage of Imood-positive cells in CD4+ and CD8+ T cell populations, comparing control and stimulated conditions within individual donors. Results are expressed as mean ± SEM (n=7 biological replicates), *P ≤ 0.05, **P ≤ 0.01, ***P ≤ 0.001, using a paired t-test. (**B**) Intracellular Imood Expression: Analysis of intracellular Imood expression was conducted following cell fixation and permeabilization, using the same antibody pairs. Contour plots display CD4+ and CD8+ T cells stained with 1B10 (blue) or 1C4 (red), with respective gates showing the percentage of Imood-positive cells. Paired before-after graphs (right) demonstrate changes in intracellular Imood expression across CD4+ and CD8+ T cell populations, accounting for intra-donor variability between control and stimulated conditions. Results are expressed as mean ± SEM (n=7 biological replicates), *P ≤ 0.05, using a paired t-test.

The trends observed in intracellular Imood expression partially mirrored those observed on cell surfaces. Specifically, intracellular Imood levels detected by 1B10 were significantly elevated upon activation in both CD4+ and CD8+ T cells, with increases of 51% and 61%, respectively (P≤0.05) (**Figure 5**). However, 1C4 staining did not reveal a significant change in intracellular Imood expression in either cell type.

These data suggest the existence of dual Imood pools within T lymphocytes: an intracellular pool, likely newly synthesized post-TCR engagement detected by 1B10, and a membrane-associated pool discernible by 1C4 staining. Differential recognition by 1C4 and 1B10 is likely due to their binding to distinct Imood isoforms, which may undergo post-translational modifications such as phosphorylation induced by TCR activation which has been previously reported (75). According to this model, the surge in 1B10 intracellular staining could be correlated with the generation of nascent Imood, which replenishes the contingent of post-translationally modified 1C4-recognizable Imood translocated to the cellular surface.

### Elevated Imood expression and polymerisation in OCD patients

In a previous study, we documented a six-fold increase in Imood mRNA levels in PBMCs of individuals with OCD (28). To corroborate these findings at the protein level, we procured blood samples from a demographically varied cohort of patients with OCD (full demographic data in **Supplementary Table 2**) and compared Imood expression with that in age- and sex-matched healthy controls.

Initial assessments revealed pronounced (p<0.01) leukopenia in the OCD group (**Figure 6A**, left panel), with leukocyte counts halved (approximately 2.5 x 10^6^ cells/mL in controls vs. approximately 1.25 x 10^6^ cells/mL in OCD subjects). Subsequent enumeration of the total lymphocyte number, as well as CD4+ and CD8+ T cells, mirrored this reduction (**Figure 6A**, middle and right panels), with both subsets exhibiting a ∼50% decrease in the OCD cohort compared to healthy controls (CD4+ cells: approximately 4 x 10^5^ cells/mL in controls versus approximately 2 x 10^5^ cells/mL in OCD; CD8+ cells: approximately 2.5 x 10^5^ cells/mL in controls versus approximately 1.25 x 10^5^ cells/mL in OCD). Analysis of Imood+ cells revealed a significant increase across the OCD cohort (**Figure 6B**), with both 1B10 and 1C4 antibodies detecting higher percentages of Imood+ cells. In CD4+ T cells, the percentage of 1B10+ cells increased from approximately 15% in controls to 33% in OCD patients, while 1C4+ cells increased from approximately 15% to 40%. Similarly, in CD8+ T cells, 1B10+ cells increased from approximately 10% in controls to 25% in OCD patients, while 1C4+ cells increased from approximately 15% to 30%. Notably, the majority of patients with OCD (approximately 60%) displayed an increase in a discrete population of Imood^high^ T cells, which were also detectable in healthy control samples (data not shown).

**Figure 6.**
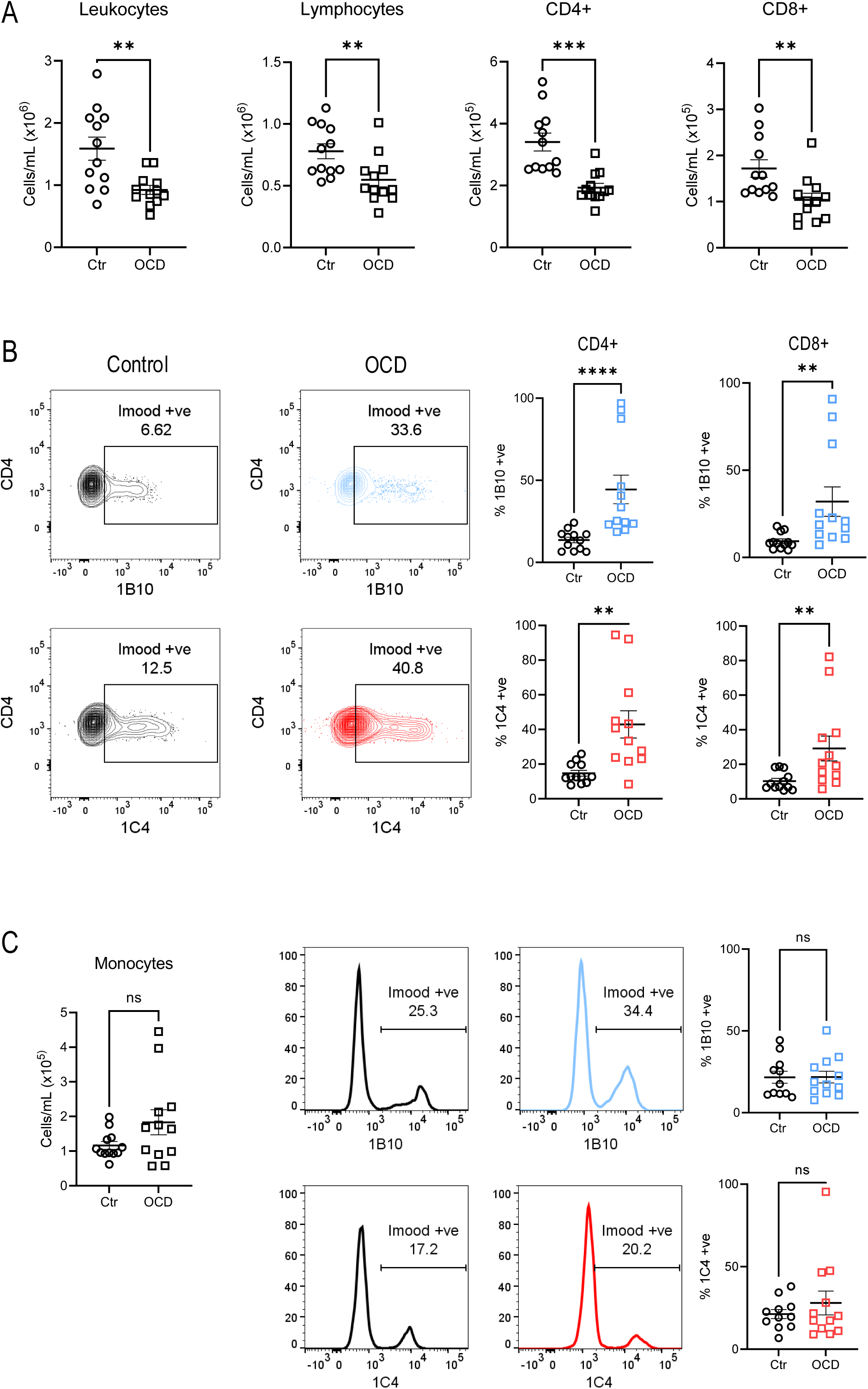
Altered leukocyte populations and increased Imood expression in OCD patients. (A) Quantification of leukocyte populations in control and OCD subjects. Scatter plots show individual data points and mean ± SEM for total leukocytes, lymphocytes, CD4+ T cells, and CD8+ T cells. OCD patients exhibit significant reductions in all cell populations compared to controls. **p < 0.01, ***p < 0.001 (statistical test not specified, presumably unpaired t-test or Mann-Whitney U test). (B) Flow cytometric analysis of Imood expression on CD4+ T cells. Left: Representative flow cytometry plots showing Imood expression detected by 1B10 and 1C4 antibodies in control and OCD samples. Right: Quantification of Imood+ cells (%) for both antibodies in CD4+ and CD8+ T cells. Bar graphs show mean ± SEM. OCD patients display significantly higher percentages of Imood+ cells for both T cell subsets and both antibodies. **p < 0.01, ****p < 0.0001 (statistical test not specified). (C) Analysis of Imood expression in monocytes. Left: Quantification of monocyte numbers in control and OCD subjects. Middle: Representative histograms showing Imood expression detected by 1B10 and 1C4 antibodies in control (black) and OCD (coloured) monocytes. Right: Quantification of Imood+ monocytes (%) for both antibodies. Bar graphs show mean ± SEM. No significant differences were observed in monocyte numbers or Imood expression between control and OCD groups. ns: not significant.

As we have previously found that PMA-or anti-CD3/CD28 activated T cells show high expression of 1C4+ on the cell surface, we wondered if the differences in surface Imood expression observed between OCD and healthy controls reflected a primed activated state of T cells. However, staining with the activation markers CD25 and CD69 showed no difference between healthy controls and patients with OCD, thus confirming a specific increase in Imood membrane expression in the T cells of these patients (**Supplementary Figure 6**). This increased expression was T cell-specific, as mononuclear cell analysis demonstrated no discernible difference between patients with OCD and healthy subjects (**Figures 6C**).

Analysis of intracellular Imood expression contrasted with the results of surface expression, as OCD T cells showed a significant reduction in both 1B10 and 1C4 MFI compared with healthy controls (**Supplementary Figure 7**). The juxtaposition of increased surface expression with a potential decrease in intracellular reserves led us to hypothesize an escalated release of Imood by T cells in the OCD population. To test this possibility, we measured Imood levels in the serum of patients with OCD and matched controls by western blotting (**Figure 7A**). As illustrated in **Supplementary Figure 8**, the aggregate serum Imood levels over the three different sets of analyses did not differ significantly between the groups. However, when examining the oligomerization state, we detected a significant decrease in monomeric Imood, coupled with an increase in trimers, in the OCD cohort relative to healthy individuals (**Figure 7B**). Cumulatively, these results delineate a profile of elevated Imood expression in T lymphocytes and altered protein polymerization in the serum of patients with OCD compared to healthy controls.

**Figure 7.**
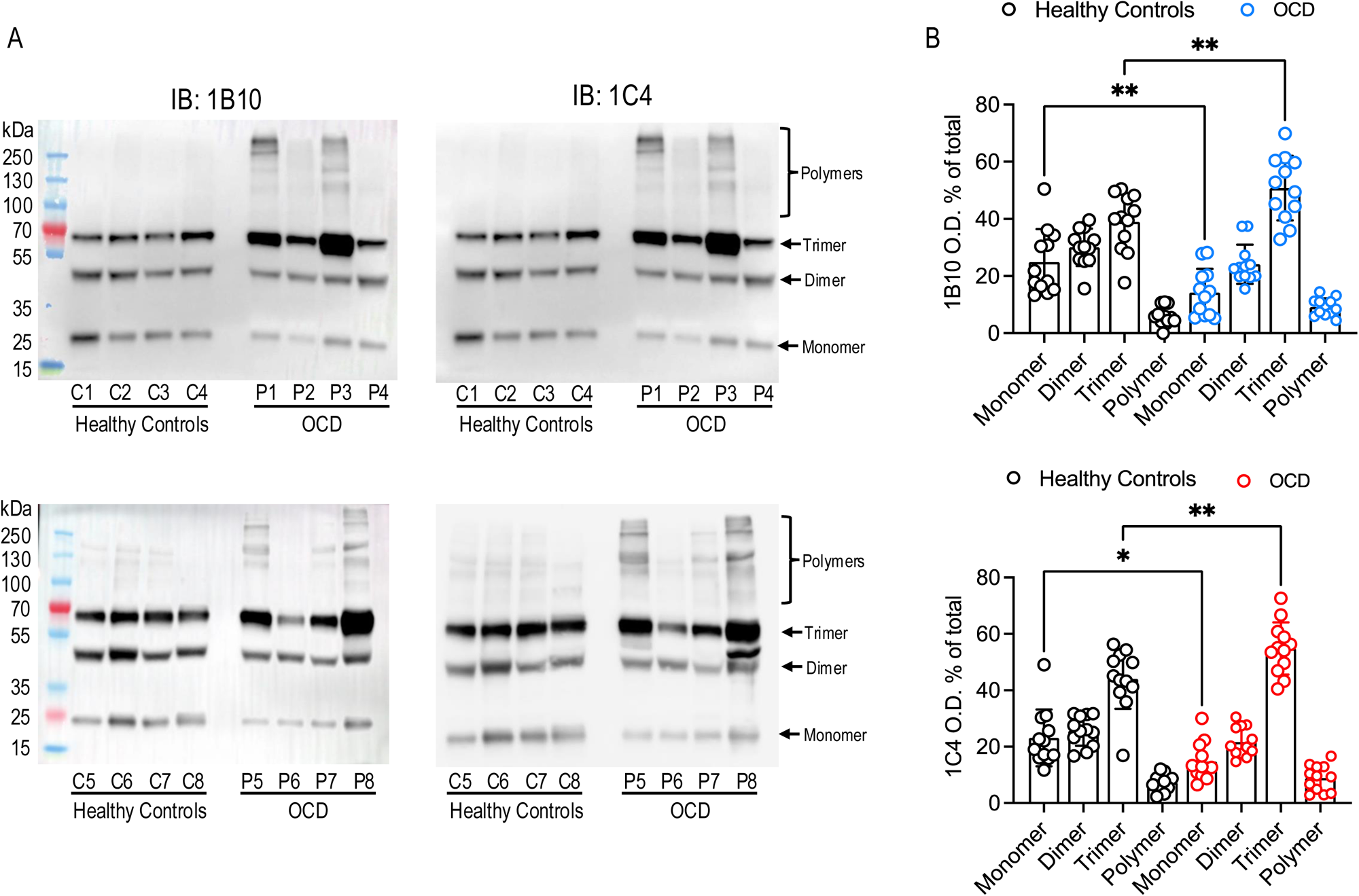
Altered Imood oligomerization state in serum of OCD patients compared to healthy controls. (**A**) Western blot analysis of Imood in serum samples from healthy controls (C1-C8) and OCD patients (P1-P8) using two different antibodies: 1B10 (top panels) and 1C4 (bottom panels). Molecular weight markers are indicated on the left. Imood oligomers are labeled as monomers, dimers, trimers, and polymers. (**B**) Quantification of Imood oligomers detected by 1B10 (top graph) and 1C4 (bottom graph) antibodies. Data are presented as optical density (O.D.) expressed as a percentage of total Imood signal. Each dot represents an individual subject. Bars indicate mean ± SEM. *p < 0.05, **p < 0.01 (statistical test not specified, presumably unpaired t-test or Mann-Whitney U test).

## Discussion

The primary objective of this study was to investigate Imood expression in circulating human immune cells. For this purpose, we used two antibodies designed to neutralize Imood *in vivo* (28). Our findings expand our understanding of the nature and biology of Imood, revealing unexpected insights into its structure, expression patterns, and potential roles in immune function and psychiatric conditions.

Our initial western blot experiments on plasma samples revealed that both 1B10 and 1C4 antibodies detect multiple forms of Imood, with bands corresponding to the dimeric, trimeric, and polymeric forms of the protein. Immunoblotting of C-Myc/DDK-tagged recombinant Imood (rec-Imood), used as a positive control, showed a higher molecular weight than expected (30 kDa vs. 24 kDa) and was detected by 1C4, but not 1B10. We attribute this increase in molecular weight to PTMs, such as phosphorylation or acetylation, which are common in recombinant proteins expressed in mammalian systems, such as HEK293T cells (76). These PTMs can also cause structural changes that affect antibody recognition, potentially explaining the differential reactivity of 1B10 and 1C4 towards rec-Imood.

To better understand the nature of the observed polymeric bands, we conducted a comprehensive bioinformatic analysis of the amino acid sequence of Imood. We found that Imood exhibits features characteristic of IDPs. Notably, both C- and N-termini were identified as the most disordered regions in the full-length protein. This finding is significant because IDRs often play crucial roles in protein-protein interactions and can drive IDP polymerization (60, 77).

Our analysis also revealed potential protein-protein interaction domains within Imood, which might facilitate self-association or interaction with other proteins, potentially contributing to oligomer formation. We identified several putative phosphorylation and acetylation sites that could explain the higher molecular weights observed for rec-Imood and the differential reactivity of our antibodies. We also found short regions with a higher likelihood of forming transient α-helices or β-strands that could serve as nucleation points for oligomerization (78, 79).

These computational predictions suggest that the bands observed in plasma samples may represent Imood oligomers. The absence of polymers in the rec-Imood samples may be explained by the fact that recombinant IDPs require mechanical agitation (shaking) *in vitro* to form oligomers and polymers that resemble those observed *in vivo* (80, 81). Although the cellular environment with its molecular crowding (82) and specific interacting partners (83) may promote oligomerization, these factors are absent in simplified *in vitro* settings (84).

Analysis of Imood expression in PBMCs indicated that in CD4+ and CD8+ T cells, CD56+ NK cells, and CD19+ B cells, Imood is primarily stored intracellularly with minimal surface expression. We observed lower expression levels of 1C4+ intracellular Imood than 1B10+, suggesting the existence of two intracellular Imood pools (unphosphorylated and phosphorylated) in equilibrium under resting conditions.

In innate immune cells, we found a similar intracellular storage of Imood in CD14+ and CD16+ monocytes, whereas CD66b+ neutrophils showed negligible Imood expression. These findings are particularly intriguing considering a previous study that identified Imood (referred to as *TDRP* in the study) as a candidate gene linked to ADHD (85). The lack of Imood in *Drosophila*, a species without an adaptive immune system, indicates a possible connection between Imood expression and adaptive immune cells.

The absence of Imood in invertebrates is particularly interesting when viewed in the larger context of immune system evolution and its unique adaptations in humans. A previous study (86) identified Imood as one of the 17 genes showing evidence of positive selection in the human lineage, indicating that these genes have undergone significant changes that became more prevalent in our species over time, potentially due to evolutionary pressures or trade-offs between beneficial and detrimental effects. Notably, all the other 16 genes identified were implicated in diseases, such as epithelial cancers, schizophrenia, autoimmune diseases, and Alzheimer’s disease.

It is interesting to hypothesize the role of Imood in behavioral responses, which, combined with its expression in the adaptive immune system, could reflect an adaptive strategy in which psychological states and immune responses are coordinated to enhance survival. This coordination could have been crucial for facing the various environmental and social challenges encountered during human evolution.

The unique behaviour of Imood in T cells, which are key players in the adaptive immune system, aligns well with the hypothesis of its possible importance in human evolution. Upon activation with PMA or anti-CD3+CD28 antibodies, T cells rapidly mobilize Imood to the cell surface. Immunofluorescence analysis revealed a polarized distribution of Imood on one side of the membrane in activated cells, reminiscent of the polarization of key early T cell signalling molecules and directional cytokine secretion.

These findings are particularly intriguing when considered alongside a phosphoproteomic study of human primary T cells that identified a phosphopeptide corresponding to the N-terminus of Imood after 5 min of TCR stimulation (75). Our cytofluorimetric data on anti-CD3/CD28-stimulated T cells showed a significant increase in cell membrane 1C4 staining, but not in 1B10 staining, whereas intracellular staining showed the opposite pattern. These findings suggest a model in which phosphorylated Imood is mobilized to the cell surface before being released into the extracellular space, whereas newly synthesized non-phosphorylated Imood replenishes the intracellular pool.

Our investigation of Imood expression in patients with OCD, previously reported to have increased Imood mRNA compared to healthy controls, supports this model. Patients with OCD showed an increase in the total percentage of T cells expressing Imood on the cell surface and higher expression levels (MFI) than healthy controls. Mirroring this increased externalization of Imood, intracellular staining showed a significant reduction in the levels of this protein in OCD samples compared with the matched controls.

We also know that the Imood promoter is differentially methylated in OCD subjects when compared to genomic material collected from healthy controls, and that this alteration is significantly correlated with the increased expression of the gene in OCD (87). We hypothesize that this increased rate of expression and possibly phosphorylation of Imood might be responsible for the higher rate of mobilization from the cytoplasm to the cell surface, where it is then released into the plasma. This extra pool of Imood produced by OCD T cells may accelerate the physiological conversion of Imood monomers to polymers.

Indeed, analysis of plasma Imood in patients with OCD by western blotting revealed an increased abundance of both 1B10+ and 1C4+ trimeric Imood compared with healthy controls, with a concomitant reduction in the monomeric form. Polymeric Imood was readily detectable in some, but not all, patients with OCD, particularly in samples with higher overall Imood expression with a marked preponderance of trimers over monomers, suggesting a correlation between the concentration of the protein in plasma and polymerization. This is in line with what has been observed for other IDPs, that is, an increase in the concentration of the monomer above a physiological threshold is one of the main factors favouring polymerization and/or liquid-liquid phase separation (88–90).

While we still do not fully understand the mechanisms underlying these differences, we speculate that there might be a correlation between disease development or severity and different scenarios of dynamic dysregulation of Imood. Nevertheless, our study offers a unique perspective on the etiopathogenesis of OCD and its related conditions. The relapsing-remitting pattern of OCD observed in some patients (91–93) could be linked to fluctuations in plasma Imood levels caused by T cell activation during infections (**Figure 8**). As T cells are activated and expand to combat an infective agent, Imood levels increase correspondingly. This surge in Imood might trigger OCD symptom exacerbation or relapse. Conversely, as the infection resolves and T cell populations contract, Imood levels decrease, potentially explaining the subsequent remission of the symptoms. Furthermore, this model might explain why therapies such as antibiotics (94–96), intravenous immunoglobulin (IVIG) (97, 98), or plasmapheresis (99–101) can sometimes lead to complete symptom remission in certain cases of OCD or PANS/PANDAS. These treatments, by modulating the immune response and potentially affecting Imood levels, could disrupt the cycle of T cell activation and symptom exacerbation, resulting in significant clinical improvement.

**Figure 8.**
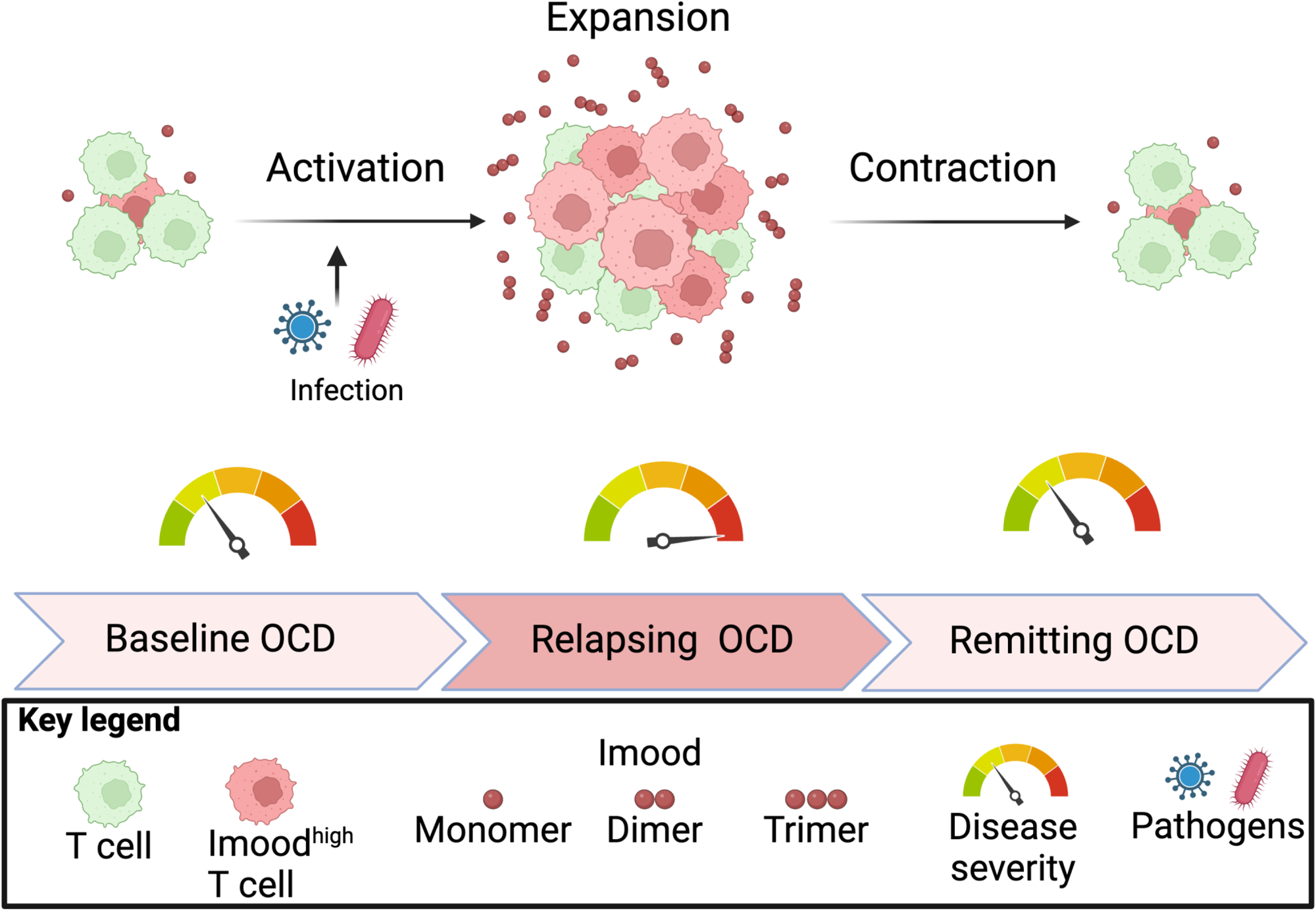
Proposed model of infection-triggered OCD relapse-remission cycles mediated by Imood dynamics. Hypothesized relationship between T cell activation, Imood levels, and OCD symptom severity. At baseline OCD state, patients maintain normal T cell populations with manageable symptoms, indicated by the lower reading on the disease severity gauge. Upon encounter with pathogens (shown as virus and bacteria), T cells undergo activation and expansion, leading to increased numbers of Imood^high^ T cells (shown in pink). During this expansion phase, elevated Imood levels are visualized by increased monomers, dimers, and trimers in the surrounding space, corresponding to the relapsing OCD state where the disease severity gauge shows heightened symptoms. As the infection resolves, T cell populations undergo contraction, returning toward baseline levels. This contraction phase is accompanied by a reduction in Imood-expressing cells and circulating Imood molecules, correlating with symptom remission as shown by the moderated reading on the disease severity gauge. Created in https://BioRender.com

In conclusion, our findings suggest that Imood may serve as a molecular bridge between the brain and the immune system. In our previous study (28), we showed that the pharmacological modulation of Imood levels with neutralizing antibodies impacts the basal anxiety levels of C57/Bl6 mice. We also acknowledge, however, that further in-depth studies will be needed to establish a causal relationship between Imood levels and/or polymerization and anxiety behavior or disease conditions. Future research should focus on elucidating the biological role of Imood in T cell function and biology as well as its potential as a biomarker for T cell activation and its specific involvement in psychiatric conditions.

All these questions need to be systematically addressed in depth, as little is currently known about Imood biology, which opens the way for further experimental and clinical investigations. We are confident that, as we continue to unravel the complexities of Imood biology, we may gain valuable insights into the intricate relationships between the immune system, brain, and mental health.

## Supporting information

Biorender publication license

## Acknowledgments

Funding: This study was supported by a CiTI and UCB Pharma funded PhD studentship to IB - Karla therapeutics (https://karla-tx.com/) and Queen Mary University of London Impact Fund – an in-kind donation of transfix tubes by Cytomark (https://www.cytomark.co.uk/) – and a crowdfunding campaign launched by OCD and PANS/PANDAS patients and families (https://www.gofundme.com/f/potential-discovery-for-ocd).

## Author contributions

IB performed all immunological analyses and screening of patient. DC and FDA designed the study, analyzed the data, wrote the manuscript, and contributed to all stages of the experimental work. MV, BdO, SF, EH and CdA recruited patients and analyzed the data. CML and MV analyzed the data and contributed to the writing of the manuscript.

## Data and materials availability

The data that support the findings of this study are available from the corresponding author upon reasonable request.

---

**Supplementary Figure 1:**
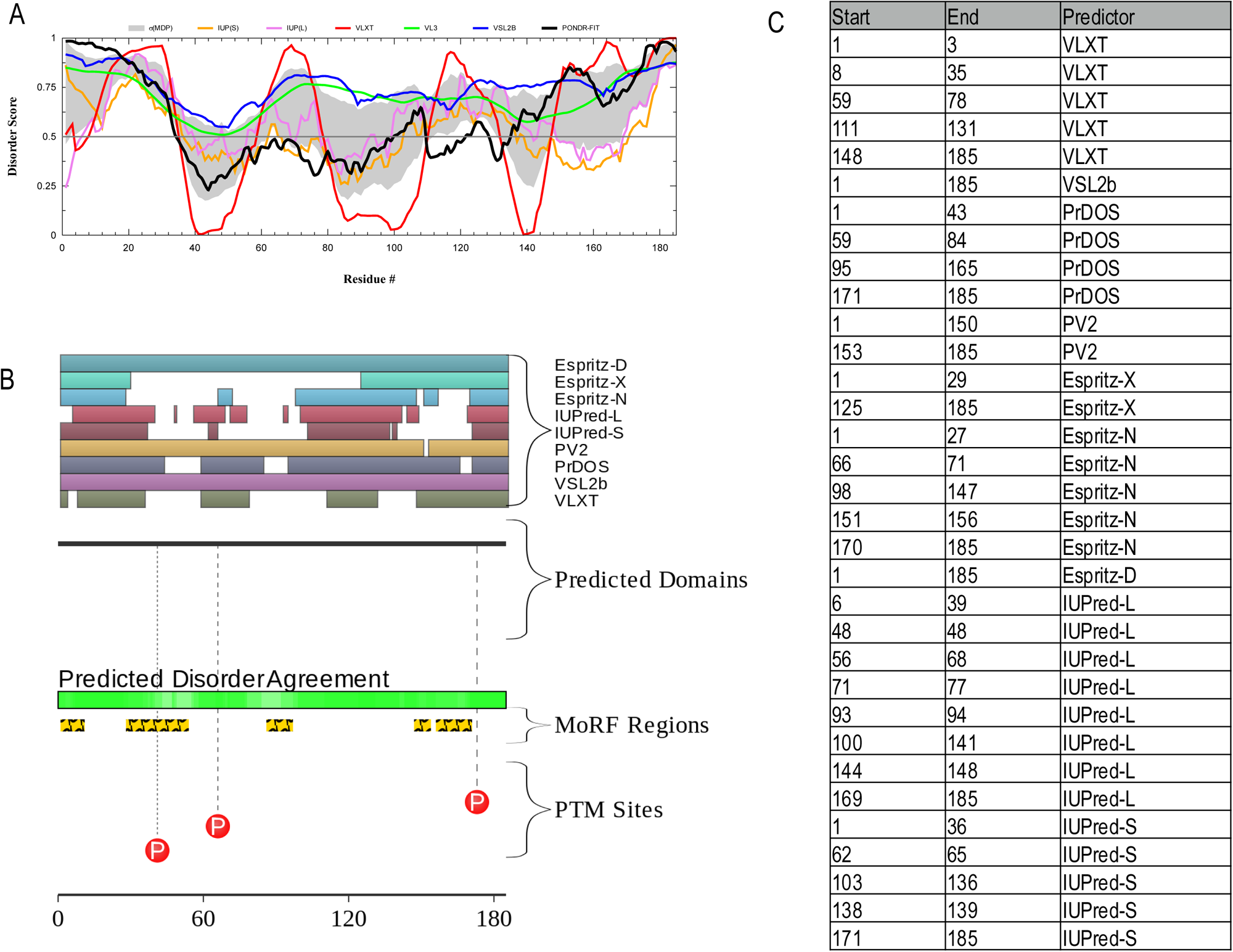
Analysis of Intrinsically Disordered Protein Regions (IDPRs) in Imood. (**A**) Disorder propensity for Imood using multiple predictors. The x-axis represents the residue number, and the y-axis shows the disorder propensity score. Different coloured lines represent various disorder prediction algorithms. (**B**) Schematic representation of predicted domains, disorder agreement, Molecular Recognition Features (MoRFs), and post-translational modification (PTM) sites in Imood. The top panel shows predicted domains using different algorithms (Espritz-D, Espritz-X, Espritz-N, IUPred-L, IUPred-S, PV2, PrDOS, VSL2b, and VLXT). The middle panel displays regions of disorder agreement among predictors (green). The bottom panel indicates MoRF regions (orange) and phosphorylation sites (P in red circles). (**C**) Detailed breakdown of predicted disordered regions, domains, and features in Imood. The table lists the start and end positions of various predicted features along with their corresponding predictors or feature types. The analysis reveals: Scattered regions of disorder throughout the protein, with strongest agreement among predictors at the N-terminus (amino acids 1-40) and C-terminus (amino acids 170-185). Five distinct Molecular Recognition Features (MoRFs) regions: 1-10, 28-53, 86-96, 147-153, and 156-170. Three predicted phosphorylation sites at Ser41, Ser66, and Ser173.

**Supplementary Figure 2:**
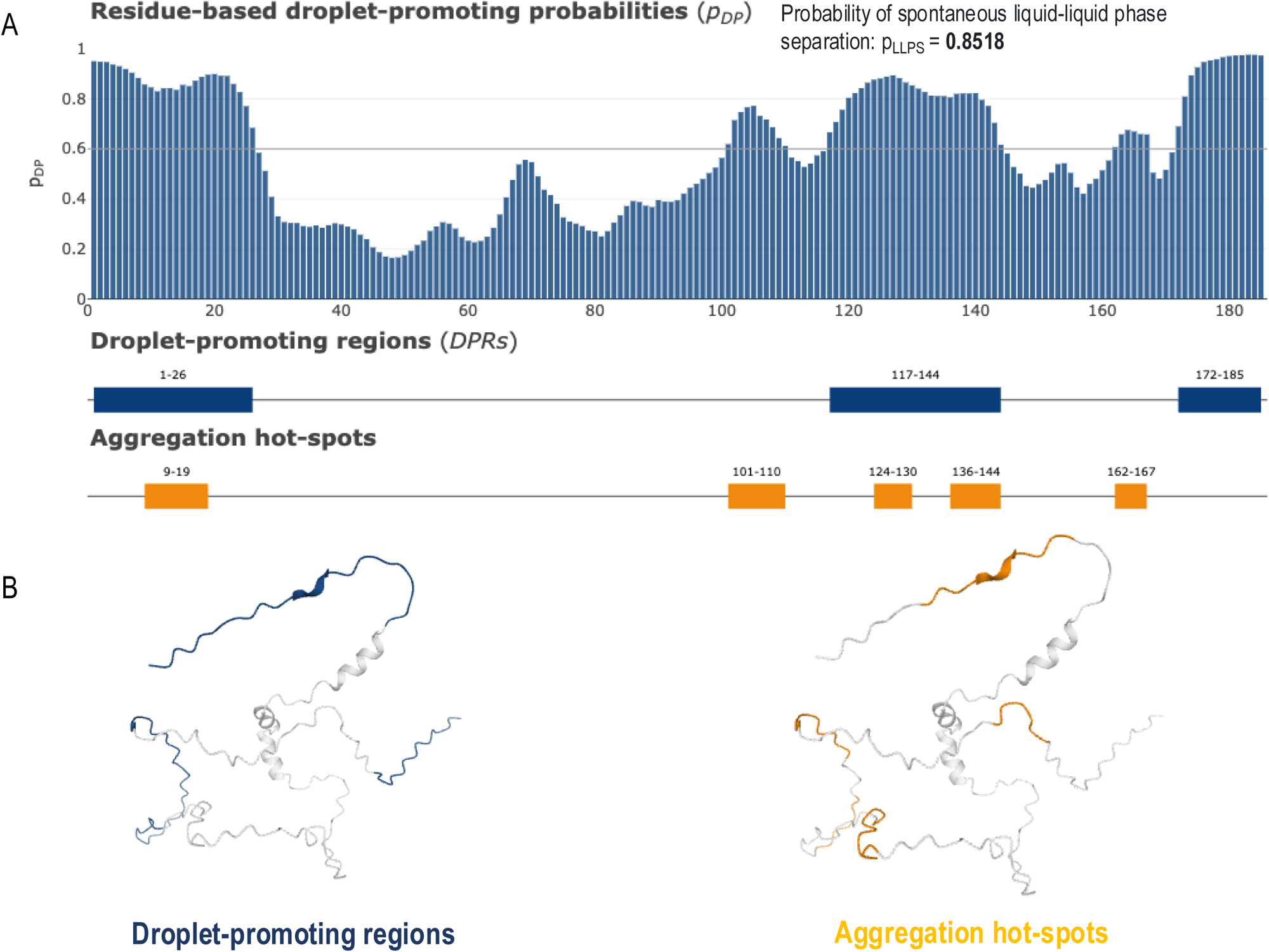
Computation analysis of the liquid-liquid phase separation (LLPS) propensity of Imood. (**A**) Residue-based droplet-promoting probabilities (pLLPS) for Imood. The bar graph depicts the pLLPS score for each residue, with the x-axis representing the residue number and the y-axis showing the pLLPS score. The overall probability of spontaneous liquid-liquid phase separation (pLLPS) for Imood is 0.85. Droplet-promoting regions (DPRs) are indicated by blue bars below the graph, spanning residues 1-26, 117-144, and 172-185. Aggregation hot-spots are shown as orange bars, corresponding to residues 9-19, 101-110, 124-130, 136-144, and 162-167. (**B**) Structural representation of Imood highlighting droplet-promoting regions in blue (left) and aggregation hot-spots in orange (right).This figure illustrates that Imood exhibits a high overall pLLPS score of 0.85, classifying it as a droplet-driving protein (pLLPS ≥ 0.60). Three main droplet-promoting regions (DPRs) are identified, along with five aggregation hot-spots. These results suggest that Imood has a strong tendency to separate into droplets spontaneously.

**Supplementary Figure 3:**
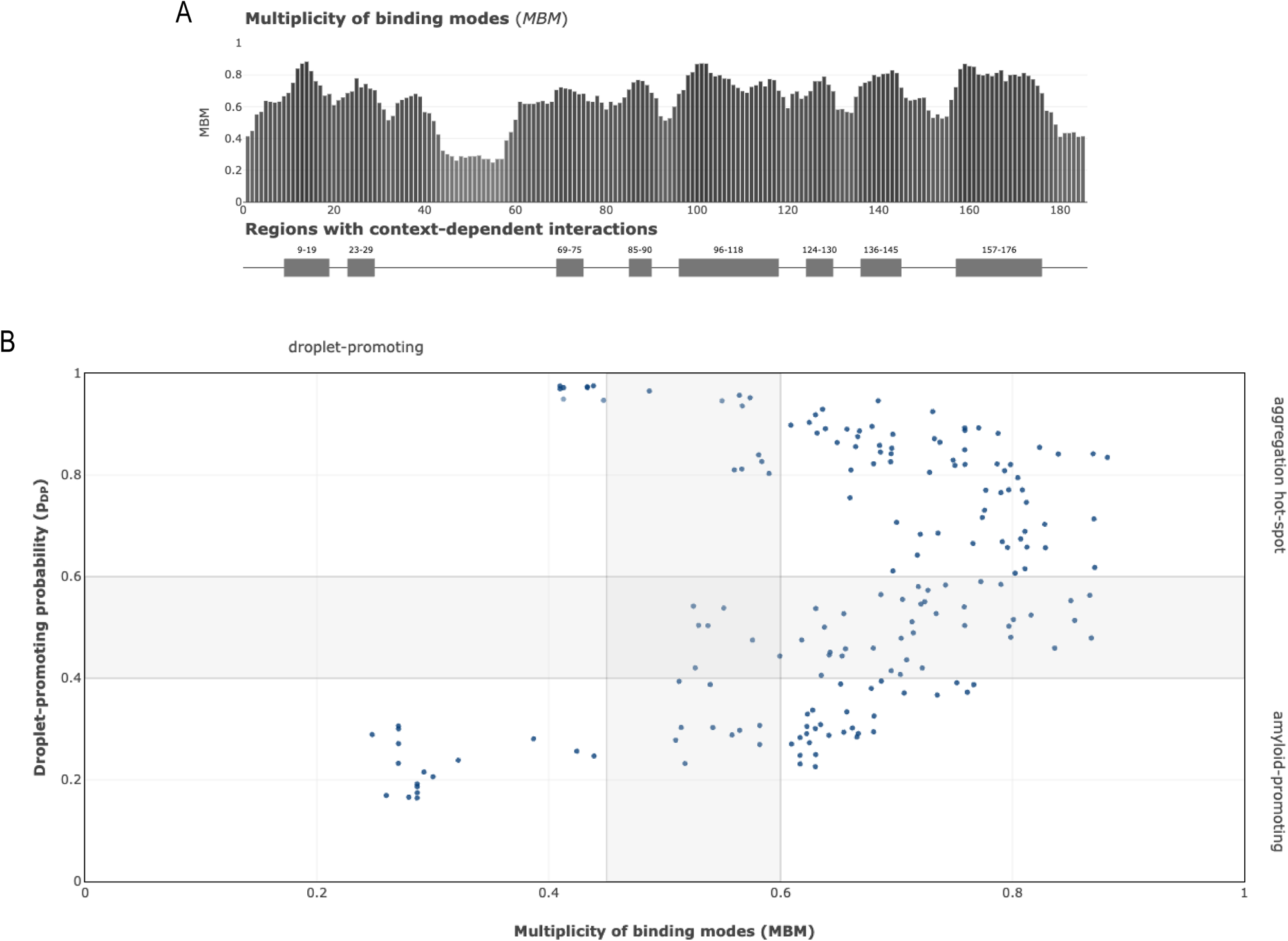
Analysis of the Multiplicity of Binding Modes (MBM) and Context-Dependent Interactions of Imood. (**A**) The upper panel displays a bar graph representing the multiplicity of binding modes (MBM) score for each residue of Imood. The x-axis denotes the residue number, while the y-axis shows the MBM score. The lower panel indicates regions with context-dependent interactions, represented by grey bars. These regions span residues 9–19, 23–29, 69–75, 85–90, 96–118, 124-130, 136-145, and 157-176. (**B**) Scatter plot illustrating the relationship between the Multiplicity of Binding Modes (MBM) and droplet-promoting probability for Imood. The x-axis represents the MBM score, and the y-axis shows the droplet-promoting probability. Each point on the plot corresponds to a residue in the Imood sequence. The plot is divided into quadrants, with the upper right quadrant indicating residues that are both highly droplet-promoting and have a high multiplicity of binding modes.

**Supplementary Figure 4:**
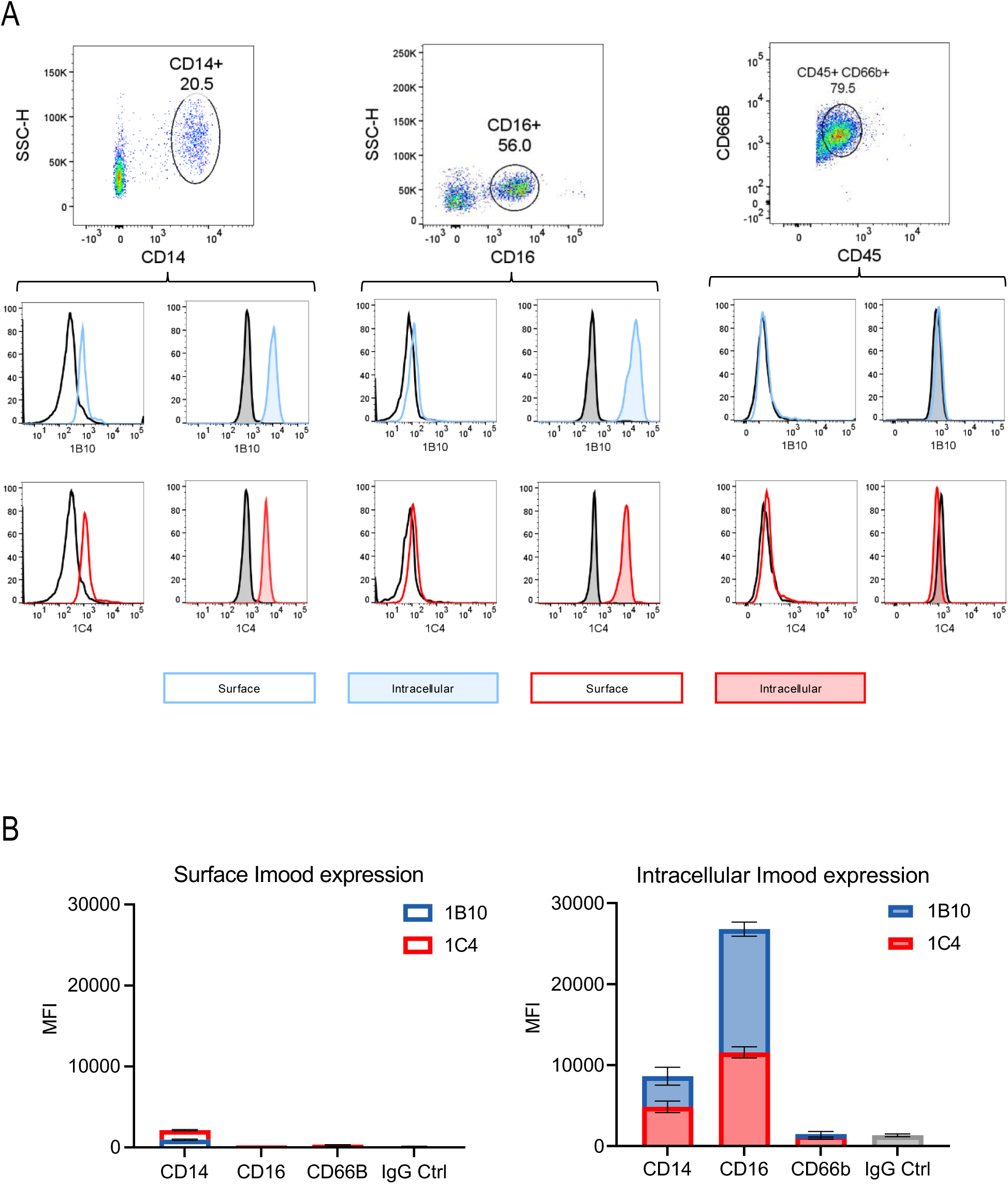
Imood expression in innate immune cells. Expression of Imood in various innate immune cell populations isolated from whole blood and analyzed by flow cytometry. **(A**) The top row shows flow cytometry plots for gating strategies to identify CD14+ monocytes, CD16+ monocytes, and CD66b+ neutrophils based on their cell surface phenotype markers. (**B**) The middle two rows display histograms for Imood staining in each cell subpopulation. Histograms show Imood surface staining (unfilled histograms) and intracellular staining profiles (tinted histograms) using mAbs 1B10 (blue) and 1C4 (red). IgG control is shown in black. The bottom panel presents cumulative stacked bar graphs showing MFI values for surface (left) and intracellular (right) Imood expression in CD14+, CD16+, CD66b+ cells, and IgG control. Blue bars represent staining with the 1B10 antibody, and red bars represent staining with the 1C4 antibody. Results are expressed as mean ± SEM (n=3 biological replicates).

**Supplementary Figure 5:**
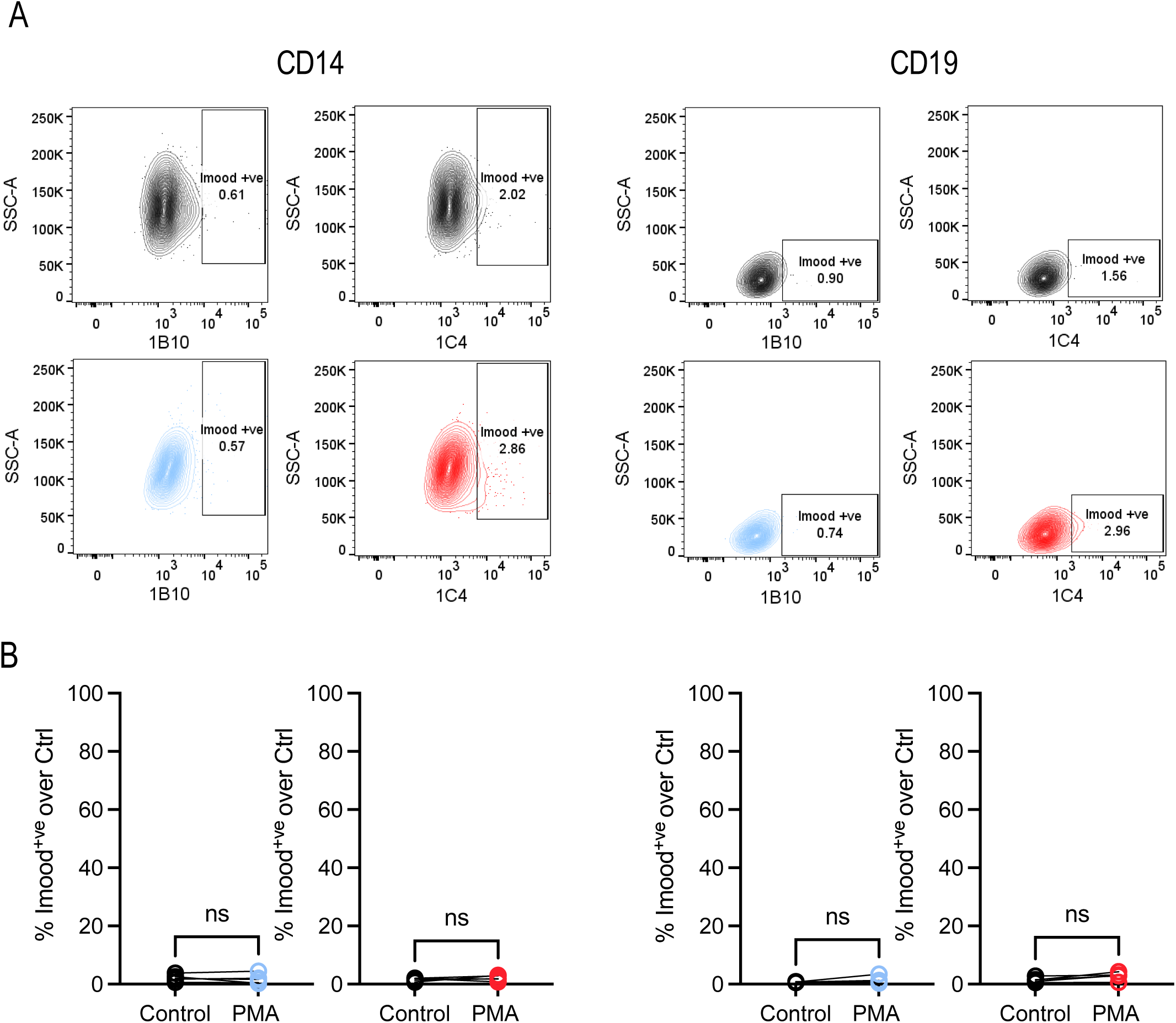
Imood expression in CD14+ monocytes and CD19+ B cells before and after PMA stimulation. **(A)** Flow cytometry plots showing Imood expression detected by 1B10 and 1C4 antibodies. For each cell type (CD14+ and CD19+), the upper panels show control conditions and lower panels show PMA-stimulated cells. Each plot includes a gate indicating the percentage of Imood+ cells. Quantification of Imood+ cells under control and PMA-stimulated conditions. Graphs show the percentage of Imood+ cells relative to control for both 1B10 (left) and 1C4 (right) staining in CD14+ monocytes and CD19+ B cells. No significant differences were observed between control and PMA-stimulated conditions (ns = not significant).

**Supplementary Figure 6.**
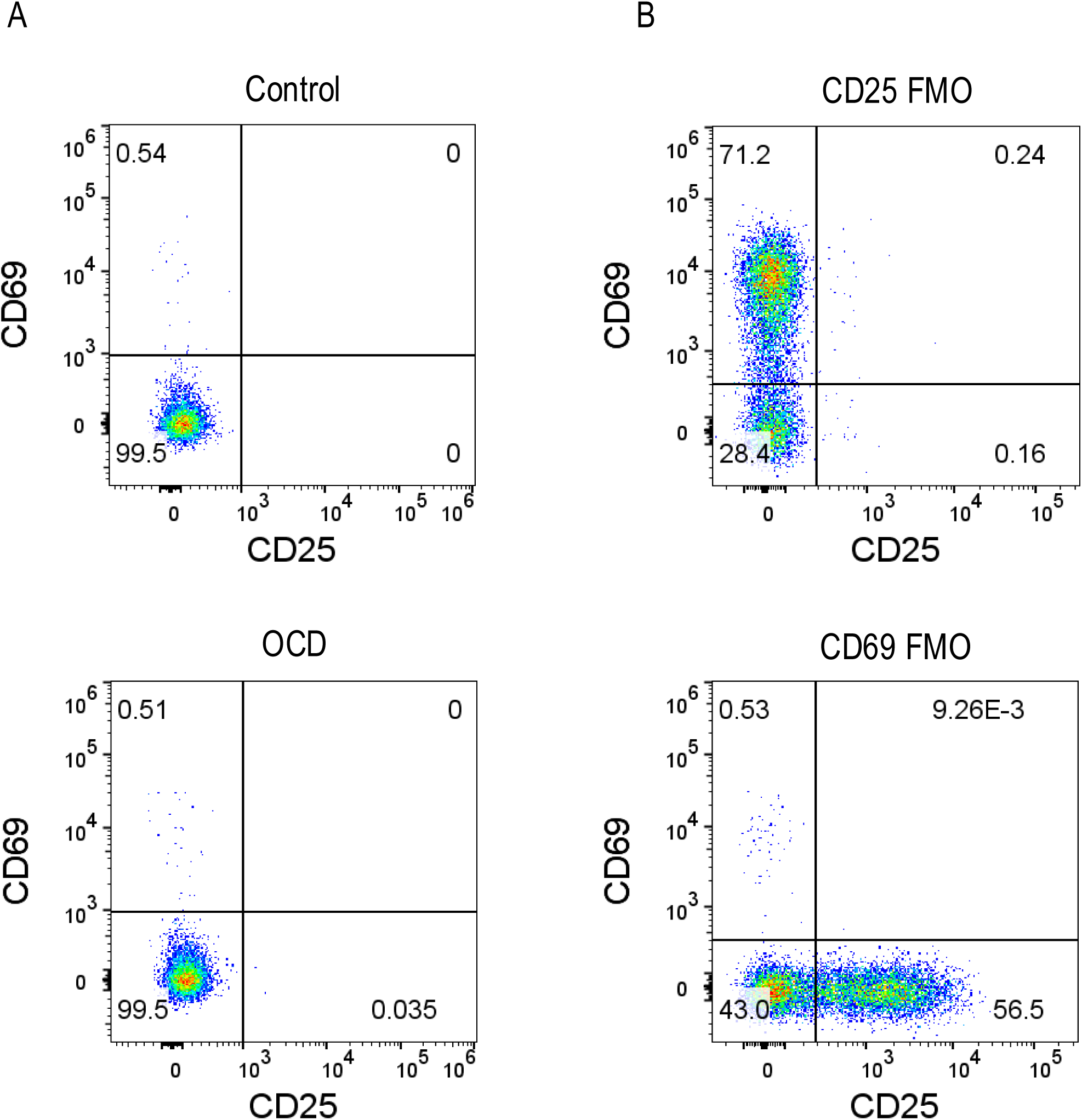
Flow cytometric analysis of CD25 and CD69 expression in T cells from control and OCD subjects. (**A**) Representative flow cytometry plots showing CD25 and CD69 expression in T cells from a healthy control (top) and an OCD patient (bottom). The percentages in each quadrant indicate the proportion of cells expressing different combinations of CD25 and CD69. (**B**) Fluorescence minus one (FMO) controls for CD25 (top) and CD69 (bottom) staining. These controls are used to set accurate gating boundaries for each marker.

**Supplementary Figure 7.**
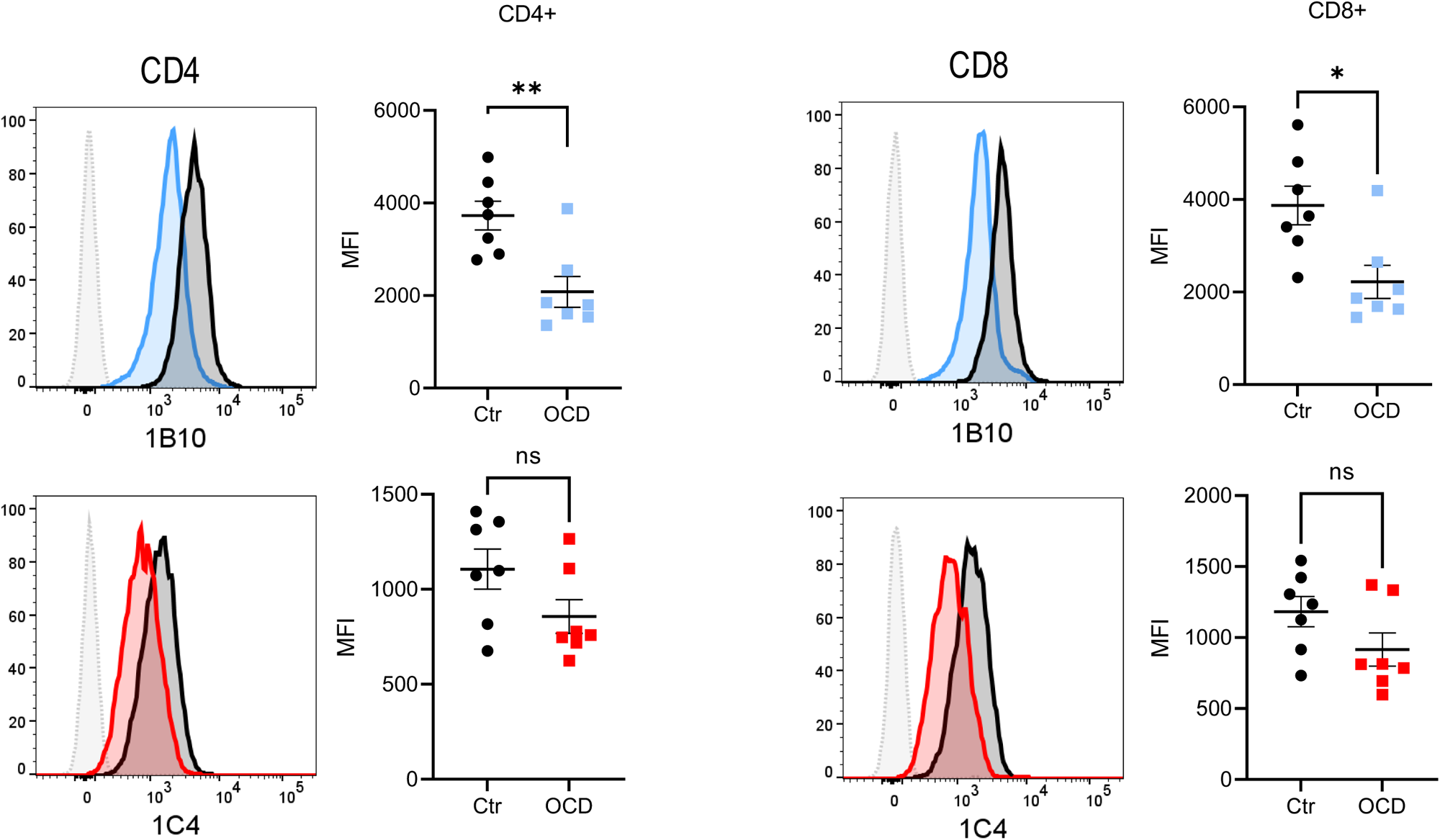
Intracellular Imood expression is reduced in CD4+ and CD8+ T cells from OCD patients compared to healthy controls. (**A**) CD4+ T cells: Left: Representative histograms showing intracellular Imood expression detected by 1B10 (top) and 1C4 (bottom) antibodies in control (black) and OCD (coloured) CD4+ T cells. Right: Quantification of median fluorescence intensity (MFI) for 1B10 (top) and 1C4 (bottom) staining. Each dot represents an individual subject. Bars indicate mean ± SEM. (**B**) CD8+ T cells: Left: Representative histograms showing intracellular Imood expression detected by 1B10 (top) and 1C4 (bottom) antibodies in control (black) and OCD (coloured) CD8+ T cells. Right: Quantification of MFI for 1B10 (top) and 1C4 (bottom) staining. Each dot represents an individual subject. Bars indicate mean ± SEM. *p < 0.05, **p < 0.01, ns: not significant (statistical test not specified, presumably unpaired t-test or Mann-Whitney U test).

**Supplementary Figure 8.**
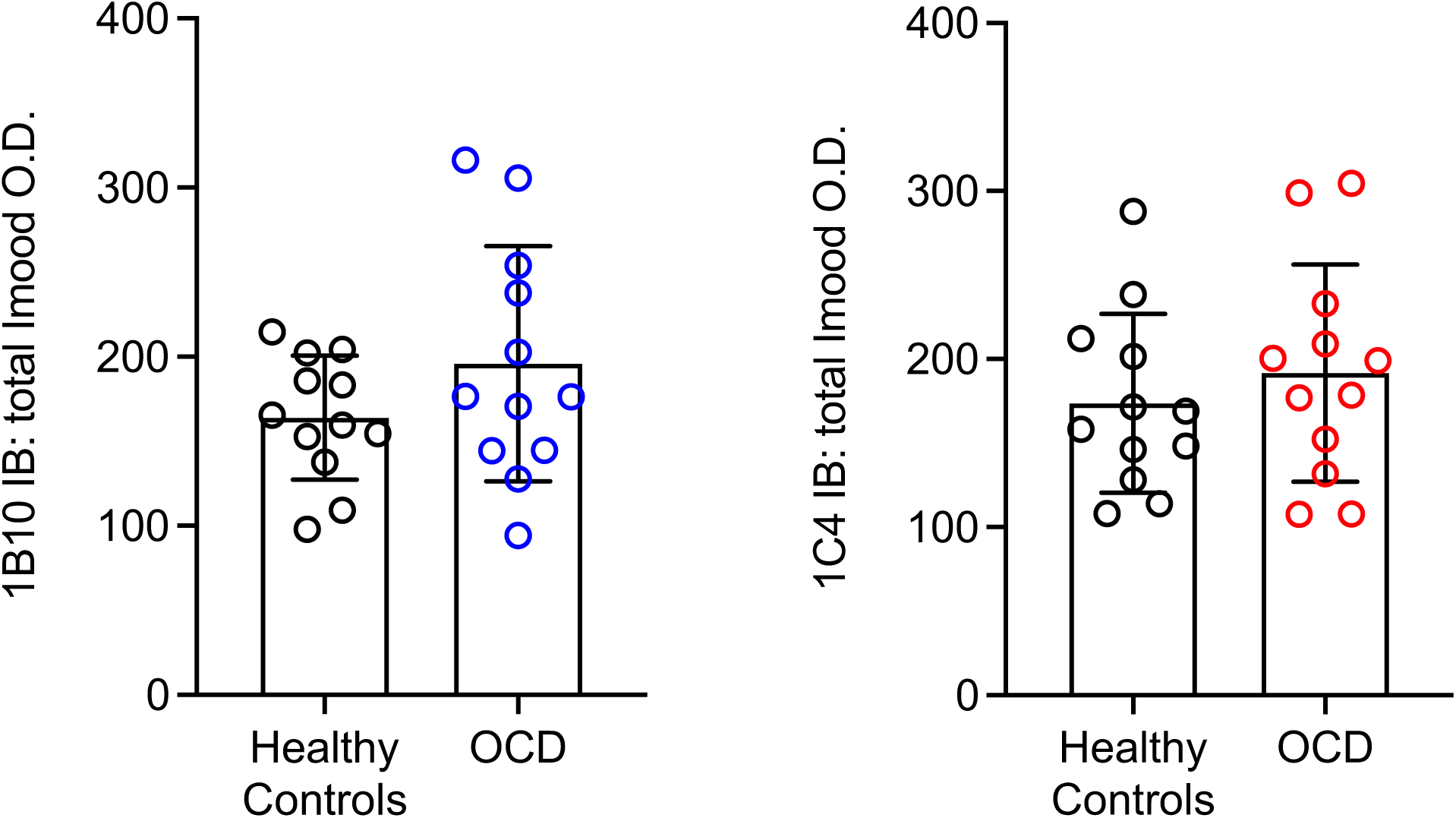
Comparison of total serum Imood levels between OCD patients and healthy controls. (**A**) Quantification of total Imood levels in serum samples from healthy controls and OCD patients, as detected by the 1B10 antibody. Data are presented as optical density (O.D.) measurements. (**B**) Quantification of total Imood levels in serum samples from healthy controls and OCD patients, as detected by the 1C4 antibody. Data are presented as optical density (O.D.) measurements. Each dot represents an individual subject. Bars indicate mean ± SD. n = 12 for each group.

