## Supplementary Table 1 for "Immuno-moodulin is Differentially Expressed in T Cells and Plasma in Obsessive-Compulsive Disorder Patients"

| Antibody | Clone | Stock | Dilution | Species | Company | Catalog number |
|---|---|---|---|---|---|---|
| **Unconjugated Antibodies** | | | | | | |
| 1b10 (mAb;IgG2c) | n/a | 2mg/mL | 1:50 | Rat | Aldevron | In-house |
| 1c4 (mAb;IgG2b) | n/a | 2mg/mL | 1:50 | Rat | Aldevron | In-house |
| IgGk2b isotype | n/a | 2mg/mL | 1:50 | Rat | R&D | MAB0061 |
| IgGk2c isotype | n/a | 2mg/mL | 1:50 | Rat | Biolegend | 400723 |
| anti-CD3 | OKT3 | 1mg/mL | 1:100 | Mouse | Biolegend | 317302 |
| anti-CD28 | 28.2 | 1mg/mL | 1:100 | Mouse | Biolegend | 302902 |
| **Conjugated Antibodies** | | | | | | |
| CD3 efluor450 | 17A2 | 5µL/test | 1:200 | Mouse | eBioscience | 48-0032-82 |
| CD4 perCP Cy5.5 | SK3 | 5µL/test | 1:200 | Mouse | Biolegend | 980810 |
| CD8 PE | HIT8a | 5µL/test | 1:200 | Mouse | Biolegend | 301064 |
| CD14 BV785 | 63D3 | 5µL/test | 1:100 | Mouse | Biolegend | 367142 |
| CD16 APC | 3G8 | 5µL/test | 1:50 | Mouse | Biolegend | 302012 |
| CD56 AF700 | HCD56 | 5µL/test | 1:200 | Mouse | Biolegend | 318316 |
| CD19 PE | 4G7 | 5µL/test | 1:200 | Mouse | Biolegend | 392506 |
| CD45 AF700 | 2D1 | 5µL/test | 1:100 | Mouse | Biolegend | 368514 |
| CD25 (APC) | BC96 | 5µL/test | 1:100 | Mouse | Biolegend | 302610 |
| CD69 (PE-Cy7) | FN50 | 5µL/test | 1:100 | Mouse | Biolegend | 310912 |
| CD11b APC | ICRF44 | 0.2µg/mL | 1:100 | Mouse | Biolegend | 301350 |
| CD66b Pacific Blue | G10F5 | 5µL/test | 1:100 | Mouse | Biolegend | 305112 |
| AF488 anti-rat | n/a | 2mg/mL | 1:200 | Donkey | Thermofisher | A-21208 |

| Antibody | Clone | Stock | Dilution | Species | Company | Catalog number |
|---|---|---|---|---|---|---|
| 1b10 (mAb) | n/a | 2mg/mL | 1:1000 | Rat | Aldevron | n/a |
| 1c4 (mAb) | n/a | 2mg/mL | 1:1000 | Rat | Aldevron | n/a |
| C8orf42 | n/a | 2mg/mL | 1:1000 | Rabbit | Novus Biologicals | NBP1-93675 |
| HRP anti-rat | n/a | 1mg/mL | 1:3000 | Goat | Abcam | ab97057 |
| HRP anti-rabbit | n/a | 2mg/mL | 1:2000 | Goat | Dako | P044801-2 |
| HRP anti-mouse | n/a | 2mg/mL | 1:2000 | Goat | Dako | P044701-2 |

| | Antibody | Clone | Stock | Dilution | Species | Company | Catalog number |
|---|---|---|---|---|---|---|---|
| | **Unconjugated primaries** | | | | | | |
| | 1b10 (mAb) | n/a | 2mg/mL | 1:50 | Rat | Aldevron | n/a |
| | 1c4 (mAb) | n/a | 2mg/mL | 1:50 | Rat | Aldevron | n/a |
| | Iba-1/AIF-1 | E4O4W | 2mg/mL | 1:100 | Rabbit mAb | CST | |
| | **Conjugated secondaries** | | | | | | |
| | AF488 anti-rat | n/a | 2mg/mL | 1:200 | Donkey | Thermofisher | A-21208 |
| | Cy3 anti-rabbit | n/a | 2mg/mL | 1:300 | Goat | | |
