## Supplementary Table 2 for "Immuno-moodulin is Differentially Expressed in T Cells and Plasma in Obsessive-Compulsive Disorder Patients"

| | |
|---|---|
| **SOCIO-DEMOGRAPHIC DATA** | |
| Gender (m;f) | 6 (50%); 6 (50%) |
| Age at recruitment (mean ± SD) | 28.8 ± 16.6 |
| Employment: | |
| graduated (%) | 58.3 |
| student (%) | 41.7 |
| Education: graduated (%) | 8.3 |
| Marital status: married (%) | 16.7 |
| **CLINICAL DATA** | |
| Age at OCD onset (years, mean ± SD) | 15.6 ± 5.2 |
| Onset <18y (%) | 50 |
| Positive family history of psychiatric disorder (%) | 50 |
| Psychiatric comorbidity (%) | 25 |
| Duration of illness (years, mean ± SD) | 13.1 ± 10.3 |
| Y-BOCS score (mean ± SD) | 17.3 ± 5.9 |
| Current treatment: | |
| antidepressants (%) | 75 |
| mood stabilizers (%) | 16.7 |
| antipsychotics (%) | 25 |
| benzodiazepines (%) | 50 |
| drug-free (%) | 16.7 |

**Legend:** Values for categorical and continuous variables are expressed in percentages and mean ± standard deviation (SD), respectively. Y-BOCS: Yale-Brown Obsessive Compulsive Scale.

**Healthy Controls** (n=12): age at recruitment (mean, SD): 30.5 ± 12.6; females 50%.
